# Beyond single discrete responses: An integrative and multidimensional analysis of behavioral dynamics assisted by Machine Learning

**DOI:** 10.1101/2021.03.17.435751

**Authors:** Alejandro Leon, Varsovia Hernandez-Eslava, Juan Lopez, Isiris Guzman, Victor Quintero, Porfirio Toledo, Martha Lorena Avendaño, Carlos Hernandez-Linares, Esteban Escamilla

**Affiliations:** Comparative Psychology Laboratory, Centro de Estudios e Investigaciones en Conocimiento y Aprendizaje Humano, Universidad Veracruzana, Xalapa, Mexico; Facultad de Estadística e Informática, Universidad Veracruzana, Xalapa, Mexico; Facultad de Matemáticas, Universidad Veracruzana, Xalapa, Mexico; Escuela de Ingeniería, Universidad Anáhuac, Xalapa, Mexico

**Keywords:** Behavioral systems, spatial-behavioral dynamics, time-based schedules, water-seeking behavior, motivational operations, Machine Learning, t-SNE, entropy

## Abstract

Behavioral systems, understanding it as an emergent system comprising the environment and organism subsystems, include spatial dynamics as a primary dimension in natural settings. Nevertheless, under the standard approaches, the experimental analysis of behavior is based on the single response paradigm and the temporal distribution of discrete responses. Thus, the continuous analysis of spatial behavioral dynamics has been a scarcely studied field. The technological advancements in computer vision have opened new methodological perspectives for the continuous sensing of spatial behavior. With the application of such advancements, recent studies suggest that there are multiple features embedded in the spatial dynamics of behavior, such as entropy, and that they are affected by programmed stimuli (e.g., schedules of reinforcement), at least, as much as features related to discrete responses. Despite the progress, the characterization of behavioral systems is still segmented, and integrated data analysis and representations between discrete responses and continuous spatial behavior are exiguous in the Experimental Analysis of Behavior. Machine Learning advancements, such as t-SNE, variable ranking, provide invaluable tools to crystallize an integrated approach for analyzing and representing multidimensional behavioral data. Under this rationale, the present work: 1) proposes a multidisciplinary approach for the integrative and multilevel analysis of behavioral systems, 2) provides sensitive behavioral measures based on spatial dynamics and helpful data representations to study behavioral systems, and 3) reveals behavioral aspects usually ignored under the standard approaches in the experimental analysis of behavior. To exemplify and evaluate our approach, the spatial dynamics embedded in phenomena relevant to behavioral science, namely *water-seeking behavior*, and *motivational opera*tions, are examined, showing aspects of behavioral systems hidden until now.

## The spatial dimension: a relevant feature neglected by regular behavioral science paradigms

The main objective of Behavioural Science is to account for the principles that underlie the Behavioural System, understanding it as an emergent and complex system comprising an environment and organism (Gibson, 1979; Kantor; 1970; Kuo, 1976; Skinner, 1938; Timberlake, 1994; Turvey, 2018). In simple words, the principal goal of Behavioural Science is to describe the principles and processes of natural behavior. The foundational works of behavioral science showed that *natural behavior* includes the Spatio-temporal dynamic as a fundamental dimension (e.g., approach-withdrawal patterns; Schneirla, 1959).

Nevertheless, for various reasons, in the history of experimental behavioral science, the temporal distribution of discrete-response analysis gained prominence over the analysis of spatial patterns and their dynamics. One reason for this was the affordable technology available at the end of the first half of the last century to make reliable and automatized behavior records and measures. These records were made primarily through the use of mechanical and electronic switches (Escobar, 2014). Thus, the methodological approach focused on computing the frequency and temporal distribution of the activation or deactivation of switches (e.g., the total number of responses to an operand, number of responses per unit of time, inter-response times, etc.). This approach was called the single-response paradigm (Henton & Iversen, 1978). Until now, it has been the standard in the experimental analysis of animal behavior (e.g., operant and pavlovian paradigms).

The predominance of apparatus, measures, data analysis, and data representations based on discrete responses (e.g., lever press, food dispenser entrance) resulted in the spatial dimension of behavior being generally neglected. It follows that standard approaches in experimental behavioral science do not account for the natural behavior associated with the organism’s movement and its embedded dynamics (León et al., 2020b).

## Computational Animal Behavior Analysis (CABA) and integration of the spatial dimension to the Experimental Analysis of Behavior (EAB)

Under operant and pavlovian paradigms, behavioral systems include complex interactions between Spatio-temporal patterns, discrete responses, and programmed stimuli, challenging to apprehend with the methodological standard approaches of the single-response record (see Henton & Iversen, 1978; Pear, 1985). Although this issue was pointed out a long time ago, it has not been easy to solve for the Experimental Analysis of Behavior (EAB). Developing an integrative approach between environmental features, discrete responses, and Spatio-temporal dynamics is still challenging.

Computational advances made in the last decade (i.e., computer vision, machine learning, and deep learning techniques) have made the recording, measurement, and analysis of spatial patterns of behavior affordable (Dell et al., 2014; Mathis & Mathis, 2020; Pérez-Escudero et al., 2014). Moreover, these technological advances to facilitate accurate and objective analysis of behavior have opened new methodological perspectives in behavioral science (Menaker et al., 2020), such as Computational Ecology and Computational Animal Behavior Analysis (CABA). Nevertheless, the EAB has so far benefited little from these developments.

Current computational methods (Datta, 2019; Mathis et al., 2018; Marshall et al., 2020; Torabi et al., 2020) provide invaluable tools to crystallize an integrative EAB approach for the analysis and understanding the Spatio-temporal dynamics (Loveless & Webb, 2021; Maekawa et al., 2020) associated with relevant behavioral phenomena (León et al., 2020a; León et al., 2020b). This multidisciplinary approach could show behavioral features and processes, hidden until now to behavioral science and, more specifically, to EAB. Hence, this emergent multidisciplinary approach could name Computational-Experimental Analysis of Behavior (CEAB).

## How the integrative approach of CEAB could extend the scope of behavioral science and Experimental Analysis of Behavior (EAB)

EAB could be positively affected by CEAB in *recording, measuring, analyzing, and representing* the behavioral systems. In addition, CEAB could help to identify features or variables embedded in the Spatio-temporal continuum of behavior under well-established methodological paradigms (e.g., operant and pavlovian conditioning), hidden until now. If it is the case, the revealed features could extend our understanding of behavioral processes and the scope of behavioral science and eventually open new research possibilities.

### Recording

The relevance of accurate and objective records to any empirical science is well known. It is established that one of the main reasons for the success of the operant paradigm was its objective record of behavior (Escobar, 2014; León et al., 2020b). Under the single-response paradigm (e.g., pressing the lever or entering the dispenser) was possible to identify ordered functional relations between different variables and procedures (e.g., schedules of reinforcement; deprivations or motivational operations) and temporal patterns of discrete responses. CEAB, recording multiple responses and spatial behavior, could close the gap to identify new interactions and determinants between environmental events and Spatio-temporal patterns of behavior.

### Measuring and data analysis

There is a strong relation between recording, measuring, and data analysis. Under the single-response paradigm, the primary measure has been the response rate (Skinner, 1966; see any current issue of *Journal of Experimental Analysis of Behavior*). The analysis has focused on unidimensional changes in this measure. The multidimensional data obtained through CEAB (e.g., through sensing of discrete responses and spatial behavior with tracking systems based on computer vision) extend measurements and analyzes coherently with an approach that assumes behavior as a Spatio-temporal continuous system (Gibson, 1979; Henton & Iversen, 1978; Kantor; 1970; Leon et al., 2020b; Pear, 1985; Timberlake, 1994). Given the vast possibilities of behavior measuring and the considerable amount of data associated with the continuous Spatio-temporal recording of behavior, the central issue is what we should measure and analyze and why (Menaker et al., 2020).

First, it is relevant to measure and analyze discrete responses (e.g., lever presses, dispenser entrances, ‘correct’ responses, among others, depending on specific behavioral phenomena) to have a comparative and parsimonious approach covering the standard paradigms in the EAB.

On the other hand, given that some very plausible proposals on the relevant functions of spatial behavior (Berlyne, 1955; De Valois, 1954; Duffy, 1957; Elliot, 1934; Henton & Iversen, 1978; Pear, 1985; Schneirla, 1959) were gradually abandoned due to the lack of record systems, due to the technology available at the time and the predominance of the single-response paradigm. It could be fertile to recover past insights about spatial behavior with current technology (Spruijt et al., 2014). The primary dimensions identified in those proposals were the *direction, intensity* (or vigor), and *variation* of behavior. The recording of both discrete responses and spatial behavior makes it possible to account for these dimensions. Thus, in second place, it could be relevant to measure and analyze the *direction*, for example, as approach-withdrawal patterns to relevant areas and stimuli (Schneirla, 1959; Duffy, 1957); *intensity*, for example, as traveled-distance, velocity, and rate of response; and *variation* of spatial behavior, for example, as recurrence patterns and entropy. Considering these dimensions in the CEAB could be a bridge to close the gap between the EAB and other paradigms of behavioral science, facilitating seeing multiple aspects of behavioral phenomena more fully. In addition, these dimensions could be helpful to identify behaviorally meaningful patterns.

### Data representation

An additional challenge is to conduct an analysis and data representation that integrates, in a perspicuous way, both discrete responses and spatial behavior as a whole behavioral system. This integrative analysis should identify and represent the participation and relevance (e.g., ranking variables) of different behavioral features or dimensions in the system (e.g., comparing the weight between variables based on discrete responses and continuous spatial behavior). Until now, this is a challenging task that could be resolved with methods for multidimensional analysis based on machine learning, such as t-SNE.

### A first approach of the CEAB

Figure 1 shows the scheme of the general procedure used in this work. Each colored row depicts a component of our approach, assisted by different computational tools and procedures, namely, *sensing/recording, measuring, data analysis*, and *representation*. Columns depict *continuous spatial behavior* and *discrete responses*, respectively. The intersection between rows and columns exemplifies some applications of computational tools, in a given component, for each kind of response (discrete or spatial).

**Figure 1.**
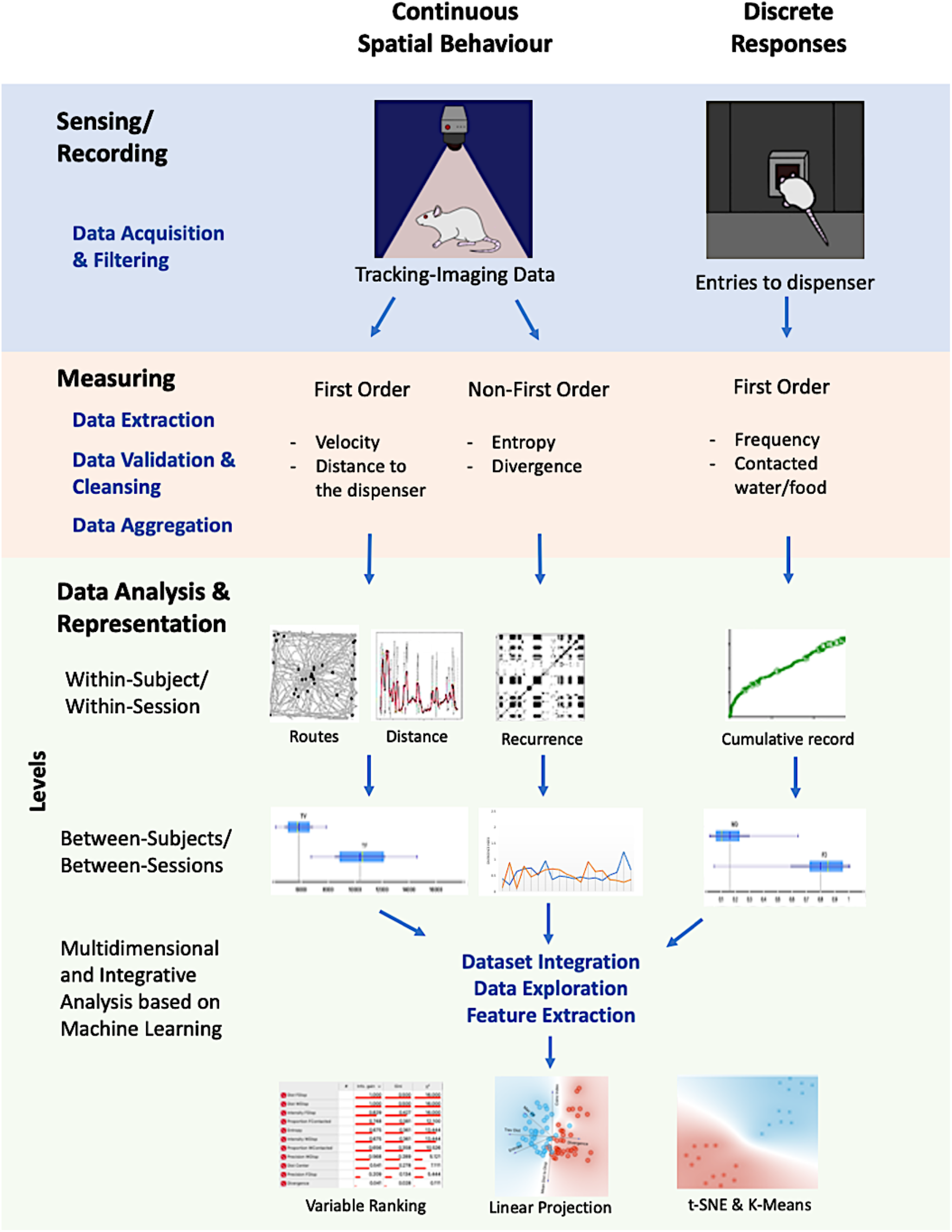
Graphical representation of the proposed integrative, multilevel, and multidimensional approach. It integrates recordings for continuous spatial behavior (based on machine vision), discrete responses of the organisms with multiple measures for each one, multilevel data analysis (within-subject, within-session, between subjects, between sessions), and a multidimensional characterization of the unitary system between Spatio-temporal dynamics and discrete responses.

The proposed approach integrates a) recording for continuous spatial behavior (based on machine vision) and discrete responses of the organisms; b) multiple measures for each record (first-order -such as *velocity, distance to focal points*- and non-first order - *entropy* and *divergence*-); c) multilevel data analysis (within-subject, within-session; between subjects, between sessions); and d) multidimensional and integrative representations of the system, made up Spatial behavior and discrete responses, based on machine learning.

Under the CEAB approach, we present two examples using a subset of datasets from our laboratory (Leon et al., 2020a; Hernandez et al., 2021) related to relevant behavioral phenomena: a) *water-seeking behavior under temporal schedules* and b) *motivational effects of water and food deprivation*. These phenomena have well-documented dynamics of discrete responses but scarcely findings concerning spatial dynamics, and even less an integrative analysis between discrete responses and spatial behavior patterns.

The purposes of the present work are a) to provide behavioral measures based on spatial dynamics sensitive to paradigmatic procedures with well-known effects on discrete responses (e.g., *stimulus schedules* and *motivational operations*); b) to reveal spatial behavioral features usually hidden under standard approaches based on single-discrete response paradigm; c) to illustrate helpful multidimensional representations, based on machine learning, for the integration of discrete responses and spatial behavior for a more comprehensive study of behavioral systems.

Although the general approach based on the CEAB is the same for both examples, given the experiments correspond to different phenomena, a specific justification, methods, and results are presented for each one. Finally, a general discussion related to both examples and the purposes of the work is presented. The hypotheses in this work are the following: 1) the proposed measures based on spatial behavior will be sensitive to EAB paradigmatic procedures; 2) the CEAB, assisted by Machine Learning, will reveal that spatial features are at least as relevant as behavioral features based on discrete responses; 3) the CEAB will show that discrete responses and spatial behavior integrate a whole behavioral system, even under different experimental procedures, and phenomena.

### Example 1

#### Water-seeking behavior: Behavioral dynamics under fixed and variable temporal schedules

One of the most significant contributions of the Experimental Analysis of Behaviour to comprehend the variables underlying behavioral phenomena are stimuli schedules (e.g., schedules of reinforcement; Ferster & Skinner, 1957). A stimuli schedule is a rule, defined in a systematic and parametric way, to present stimuli (e.g., water, food; Reynolds, 1975). The stimuli schedules can be categorized according to several criteria; one of the most common and useful is contingent vs. non-contingent schedules. The first one, in methodological terms, is usually associated with operant contingencies and the second one with pavlovian contingencies. The difference between contingent and non-contingent schedules is that in the first one, the occurrence of a given stimulus is dependent (or contingent) on a given response of the organism (e.g., lever presses). In contrast, in non-contingent schedules, the presentation of the stimuli does not depend on any organism’s response but only on the temporal relation between the stimuli. These last are named time-based schedules.

There is a vast corpus of research, with both kinds of schedules, with several species and apparatus (Boren et al., 1978; Lachter et al., 1971; León et al., 2020a; Hernandez et al., 2021; Zuriff, 1970). Most of this research is based on the recording and data analysis of a single discrete response, especially with ‘appetitive’ stimulation. The primary data are head-entries to a food or water dispenser and food pellets or drops of water consumed, in other words, the temporal distribution of a given discrete response. Only in a few studies, the data was extended to time spent in zones near a dispenser (Baum, & Rachlin, 1969). Different behavioral phenomena have been studied with time-based schedules, such as ‘superstitious behavior’ (Reberg et al., 1977; Skinner, 1948), ‘timing’ (Drew et al., 2005; Sanabria et al., 2009), among others. These different phenomena could be characterized as behavioral systems and their corresponding spatial-temporal dynamics from a systemic and parametric approach. The effects of the two time-based schedules, fixed and variable, on behavior have been scarcely studied comparatively. On the other hand, no comparative studies explicitly included the spatial dimension of behavior and its dynamics. Under these fixed and variable temporal schedules, neither integrates the standard data based on discrete responses with continuous data based on locomotion.

Under this rationale, we evaluated the spatial dynamics of behavior in Wistar rats under two temporally water-delivery schedules (fixed and variable-time schedules) in a Modified Open Field System (MOFS) from a multidimensional analysis, using machine learning tools.

## Method

### Subjects

Four experimentally-naïve female Wistar rats were used, two rats were assigned to a Fixed-Schedule Condition and two rats to a Variable-Schedule Condition. All rats were three months old at the beginning of the experiment. Rats were housed individually with a 12-hr light and dark cycle and maintained under a daily schedule of 23 hours of water deprivation with free access to water 1 hr. after experimental sessions. Food was freely available in their home cages. One session was conducted daily, seven days a week. All procedures were conducted according to university regulations of animal use and care and followed the official Mexican norm NOM-062-ZOO-1999 for Technical Specification for Production, Use, and Care of Laboratory Animals.

### Apparatus

A Modified Open Field System (Model WEOF by Walden Modular Equipment) was used. A diagram of the apparatus can be found in León et al. (2020a). Dimensions of the chamber were 100 cm x 100 cm. All four walls of the chamber and the floor were made of black Plexiglas panels. A water dispenser (by Walden Modular Equipment), based in a servo system, when activated, delivered 0.1cc of water on a water cup that protruded from the center of the MOFS. The MOFS was illuminated by two low-intensity lights (3 watts) located above the chamber and on opposite sides of the room to avoid shadowed zones. Once delivered, the water remained available for 3 s. A texturized black patch, 9×9 cm with 16 dots/cm, printed in a 3d printer, was located close to the water dispenser to facilitate its location.

The experimental chamber was located in an isolated room on top of a table of 45 cm in height. The room served to isolate external noise. All programmed events were scheduled and recorded using Walden Tracking System (v.0.1). In addition, a Logitech C920 web camera recorded rats’ movement at the center, located at the center, 1.80m above the experimental chamber. Tracking data was analyzed using Walden Tracking System (v.0.1). This software recorded rats’ location, by the center of mass, every 0.2s in the experimental space using a system of X, Y coordinates. Data files obtained from this software were then analyzed using MOTUS^®^ and Orange 3.26 Software.

### Procedure

Subjects were exposed to one of both conditions of water delivery: a) Fixed Time (FT) 30 s schedule or b) Variable Time (VT) 30s schedule. Each condition lasted 20 sessions. Each session lasted 20 minutes. Rats were directly exposed to the conditions without any previous training. Two rats were assigned to Condition 1 (FT) and two to Condition 2 (VT).

### Data analysis

To have a complete representation of the behavioral system, we analyzed different dimensions and levels based on the record of spatial behavior in a bi-dimensional space at 5 frames/s; these are described below. Formal and computational descriptions of the measurements and methods of analysis are found in the supplementary material.

#### Analysis between subjects within-session

This level of analysis was conducted, with representative subjects, thorough visual inspection of the data to identify changes in the spatial dynamics, moment to moment, related to water deliveries and the water dispenser location, through the sessions. The measures and representations used to account for the changes in *direction* and *variation* of spatial behavior under the different experimental conditions (FT vs. VT). In addition, they allow depicting the evolution and process of the spatial behavior to compare the experiment’s initial, intermediate, and final session. The specific analyses for this level were: *Bidimensional routes* and *rat’s location* at the moment of water delivery per session; *Distance to the dispenser*, moment to moment (5 frames/s), and *smoothed distance* to the dispenser with a moving average of 200 frames (for a formal description, see supplementary material); *Recurrence plot*, this plot depicts the change of regions of each rat in a matrixial configuration of 10 × 10 virtual zones (for a description, see supplementary material).

#### Analysis between subjects throughout the experiment

This level of analysis was conducted, through visual inspection, to identify the stability or variation of spatial behavior throughout the whole experiment. The used measures were *entropy* to indicate the variability of the organism’s location and *divergence* to indicate the consistency or inconsistency in such variability between consecutive sessions (for a formal description of *entropy* and *divergence*, see supplementary material).

#### Analysis between conditions by feature for all sessions

This analysis and data representation level was conducted to identify the experimental condition’s global effect on each spatial and discrete feature (FT vs. VT). The representation and analysis were based on measures of central tendency and variance. The analyzed features were: *traveled distance, entropy, divergence, maximum velocity, coincidence index, mean distance to the dispenser*. These features account for *intensity, direction*, and *variation* of behavior.

#### Multidimensional and integrative analysis based on Machine Learning

The main level of analysis allows the integration of a complete comparison and representation of all features in a whole behavioral system. *Ranking variables, t-SNE*, and *linear projection* were conducted (for a formal description, see supplementary material) to identify the weight of each feature and the effect of the experimental condition on the multidimensional system as a whole. Data of all subjects and sessions were used.

## Results

We conducted an integrative and multilevel analysis to characterize the behavioral continuum and compare the spatial dynamics of behavior under fixed and variable time schedules. First, we present representative within-subject results of the behavioral continuum within sessions, for the first, intermediate, and last session of the experiment, for one representative subject for each condition. Second, we show summary results for measures based on spatial behavior of first and non-first order (entropy and divergence) and a measure based on discrete responses (coincidence index). Third, we present a multidimensional analysis and integrative representation based on Machine Learning.

Figure 2 shows routes (Panel A), relative distance to the dispenser (Panel B), and recurrence plots (Panel C) for one rat under Fixed Time Schedule (Rat 1, left section) and one rat under Variable Time Schedule (Rat 4, right section). Each column corresponds to a given session (1, 10, and 20). In Panel A, in a bi-dimensional representation of the MOFS, the routes of the rat (grey lines) for the whole session and the rat’s location at the moment of water delivery (Location at Water Delivery, LWD) are presented (black dots). Three findings are worth mentioning: 1) routes were more extended under Fixed Time (FT) than with Variable Time (VT); 2) for both conditions, a progressive change in the direction of the routes was observed, toward the water-delivery zone, as the sessions progressed; and 3) the LWD gradually got closer to the water dispenser as the experimental sessions progressed. Panel B shows the relative value of the distance from the rat to the dispenser every 0.2 s (grey dots). In addition, to show the tendency of the distance function, we performed a smoothing of it (red line) by using a moving average of 200 frames (i.e., 40 seconds, see equation in the supplementary material). Values close to 1 indicate that the rat’s distance to the dispenser was the maximum possible; values close to zero indicate that the rat was located in a close location to the dispenser in a given time (frame). Under both conditions, a back-and-forth pattern was observed (grey dots), but this was more pronounced and had shorter periods under FT than in VT. In addition, the moving average (red line) suggests a tendency, under both conditions, to reduce the distance to the dispenser as the experiment progresses.

**Figure 2.**
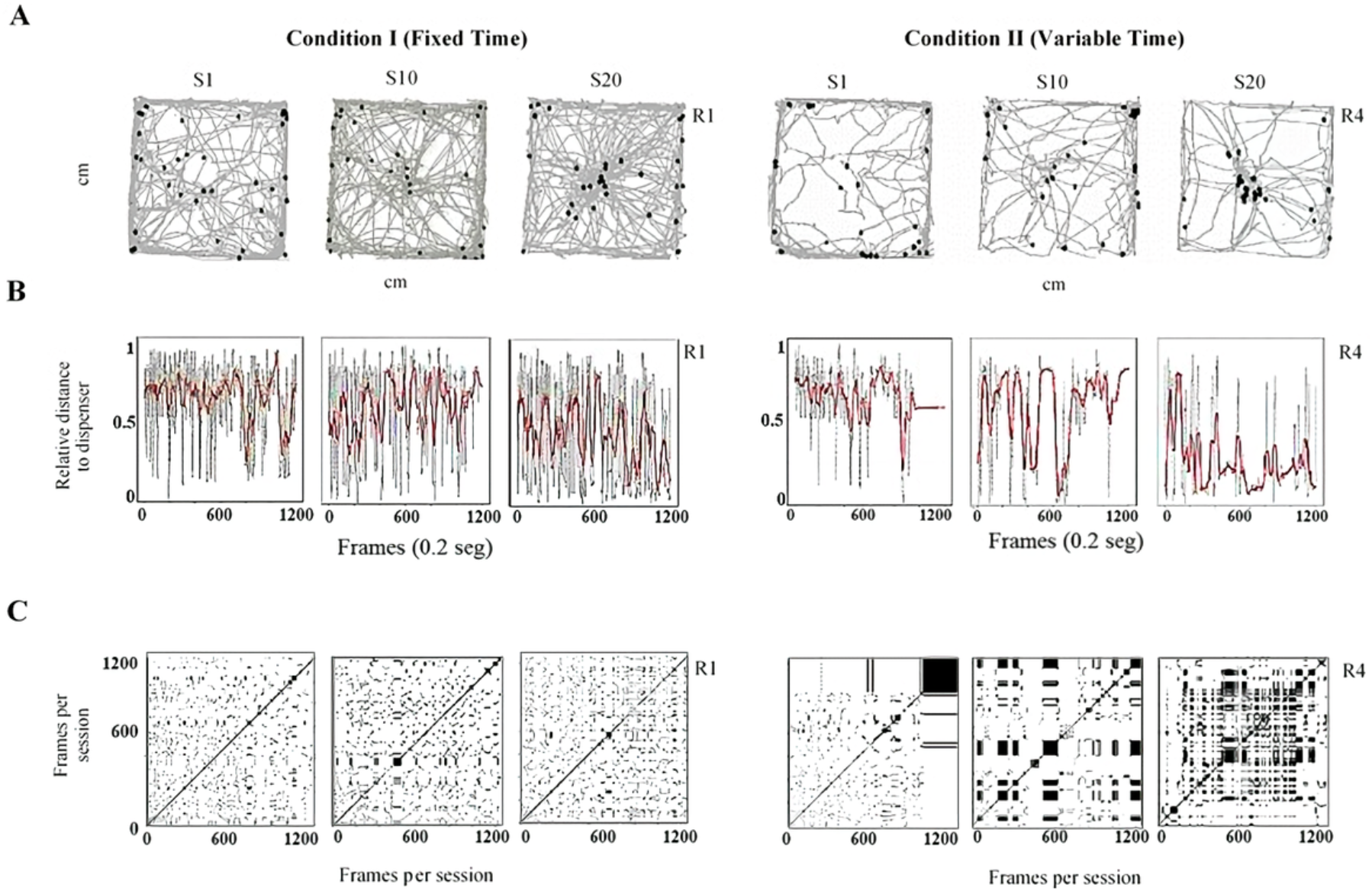
Representative within-subject results of the behavioral continuum for the initial (Session 1), intermediate (Session 10), and last session (Session 20) for one rat under Fixed Time (R1, left panels) and one rat under Variable Time (R4, right panels). Each column depicts a session. Panel A shows the routes of the rat (grey lines) and the rat’s location at the moment of water delivery (black dots) in a bi-dimensional representation of the experimental chamber. Panel B depicts the relative distance to the dispenser (Y-axis) moment-to-moment (X-axis). Finally, panel C shows recurrence plots. See the text for a complete description.

Finally, Panel C shows recurrence plots. This plot depicts the change of regions of each rat (in a matrixial configuration of 10 × 10 virtual zones) as the session progresses (see supplementary material). Both axes show time on a time frame of 0.2 s. If a rat was on a *R*_*k*_ region in a *T* time and *T* + *n* was in the same region; a black mark represented the recurrence in a given location. On the contrary, on *T* + *n* the rat was on a different location, a white mark would be shown. The densification and alternation of black-white checker patterns indicate high recurrence to a given region; a higher proportion of continuing black zones would mean higher permanence. A higher proportion of white zones would mean extended transitions among regions. Panel C shows a perspicuous difference between both conditions, higher recurrence under FT than in TV, and higher permanence in zones under VT than FT. Figure 3 shows the entropy (Panel A) and divergence (Panel B) values per session for each rat for both conditions. The entropy is helpful in the context of this experiment as a measure of the variation of locomotion patterns and the dynamic of behavior (see methodological and mathematical description in the supplementary material). For our subject matter, higher entropy represents high variation and dynamics of spatial behavior. In Panel A, the similarity between entropy plots within the condition and the difference between conditions are clear. The entropy was higher under FT than VT. A divergence index was calculated to determine the variations of spatial behavior between consecutive sessions (Panel B). This index was calculated by comparing the distribution of the organism’s locomotion into the arena between two consecutive sessions (e.g., 1 and 2; 2 and 3, etc.). A value close to zero indicates no difference in the distribution of locomotion between sessions; a value far from zero indicates a difference in the distribution of locomotion between complete sessions (see mathematical description in supplementary material). Panel B shows that the divergence was lower and more stable under FT than VT; this implied more variation of spatial behavior between consecutive sessions under VT.

**Figure 3.**
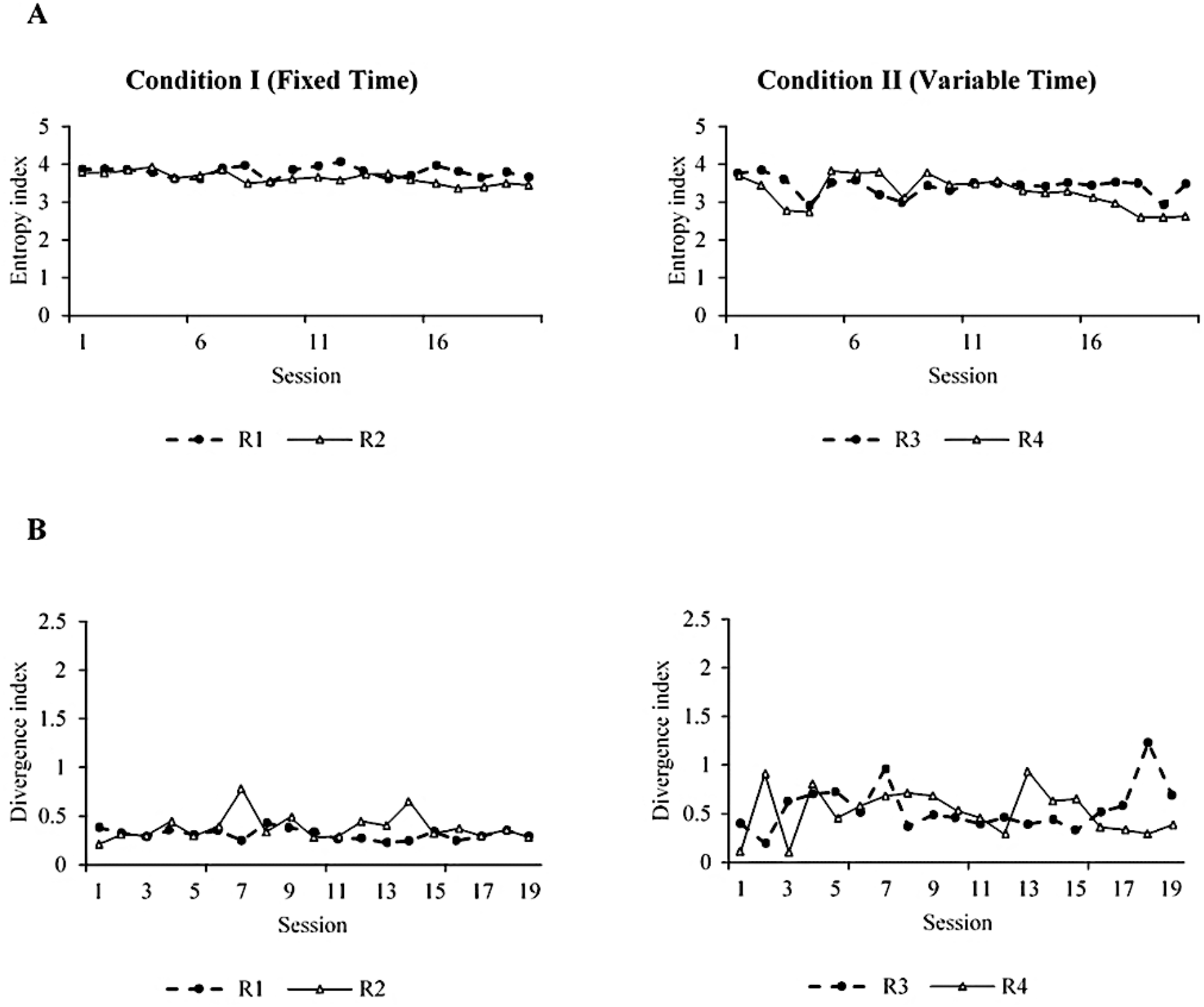
Entropy (Panel A) and Divergence (Panel B) measures for subjects R1 and R2 under Fixed Time schedules (left) and subjects R3 and R4 under Variable Time (right).

Figure 4 shows summary results of all sessions and subjects related to the spatial dimension of behavior. The traveled distance per session (Panel A), entropy (Panel B), and maximum velocity in a given frame per session (Panel D) were higher under FT than VT. In contrast, the divergence (Panel C) was significantly higher under VT than FT. On the other hand, the mean distance to the water dispenser per session (Panel F) was more dispersed under VT than TF and slightly higher, but the difference between both conditions was not robust. Finally, the coincidence index (Panel E) of the location of the organism in the dispenser zone (10 cm radio around to the dispenser allocation) at the time when water was available (3 s) was higher under FT than VT. The coincidence index is relevant because it is closely related to standard paradigms based on discrete and single response recording (e.g., entries to the dispenser).

**Figure 4.**
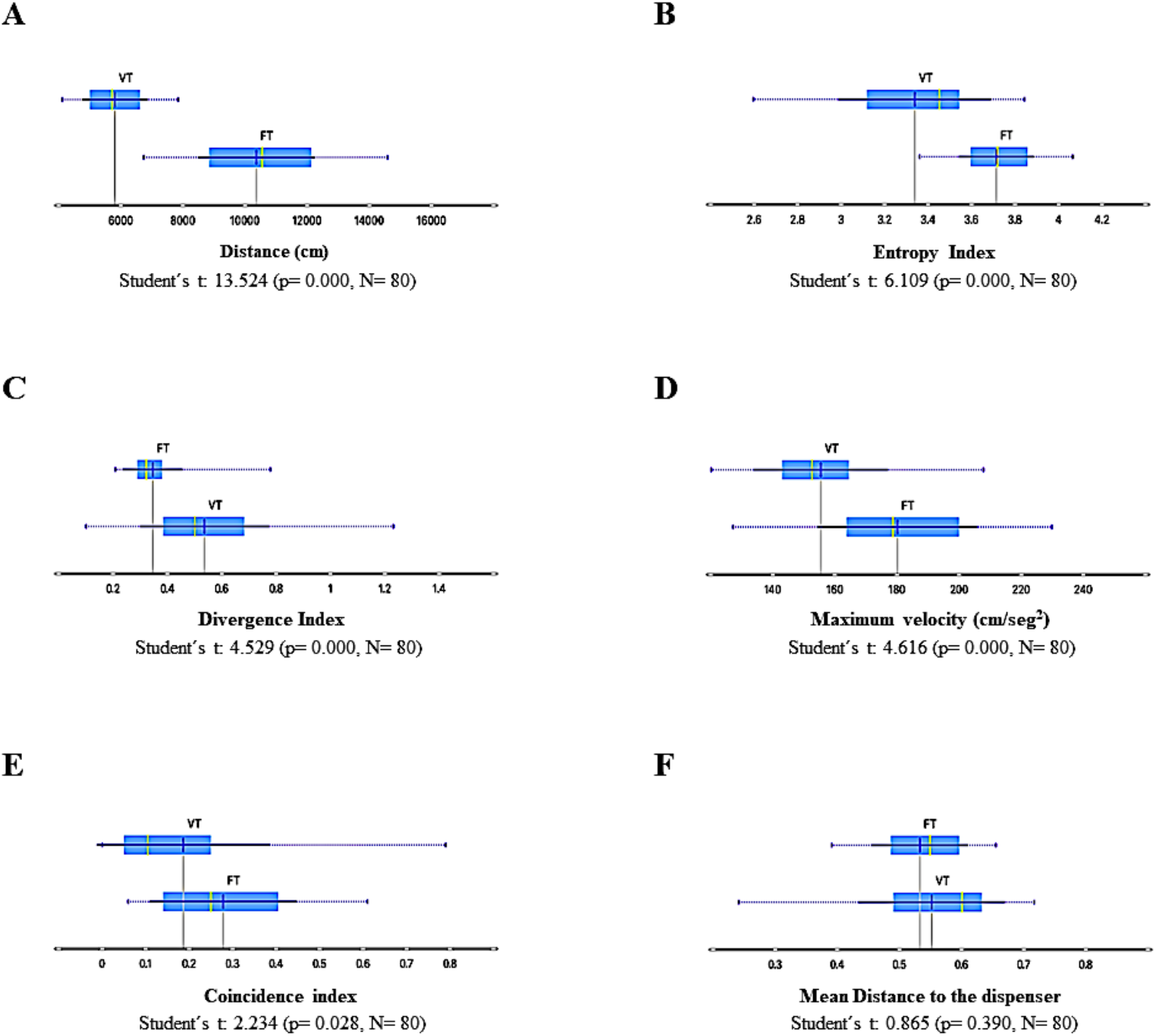
Summary results of spatial-behavior measures for all sessions and subjects under Fixed Time (FT) and Variable Time (VT) schedules. Panel A, traveled distance per session; Panel B, entropy; Panel C divergence; Panel D, maximum velocity per session; Panel E, coincidence Index; and Panel F mean distance to the dispenser. Each box depicts the mean (dark blue vertical line), the median (yellow vertical line), the standard deviation (thin blue line), and the values between the first and the third quartile (blue highlighted area).

To identify the relevance of each dimension or variable in the emergent behavioral system, understanding it as the functional interdependent relationship between variables concerning each condition, we conducted a *variable-ranking* analysis based on Machine Learning. The *variable ranking* consists of ordering a set of features by the value of a scoring function (measuring the relevance of each feature) given a target as a predicting tool, in our case, the experimental condition. *Variable ranking* allows knowing the importance or relevance of the features that better explain a target variable (for a complete description, see supplementary material). Specifically, we used filter algorithms, the most used given the low computing resources used for applying them even on high dimensional datasets. Given our subject matter, the integrative analysis of spatial dynamics of behavior with discrete responses under two different time-based schedules (FT vs. VT) and our datasets, we applied three theoretical information filter algorithms for single *variable-ranking*, namely *information gain, mean decrease impurity Gini index* and χ^2^ (for a complete description concerning these algorithms, see supplementary material), for *coincidence Index, mean distance to the dispenser, traveled distance, maximum velocity, entropy*, and *divergence*.

Figure 5 shows that according to the *variable-ranking* procedures, the most relevant features were related to the spatial dimension of behavior. These were *traveled distance, entropy*, and *divergence*. On the other hand, the *coincidence index*, the most closely variable related to the measures of the standard paradigms based on discrete response recording, was less relevant than the other features related to the spatial dimension of behavior.

**Figure 5.**
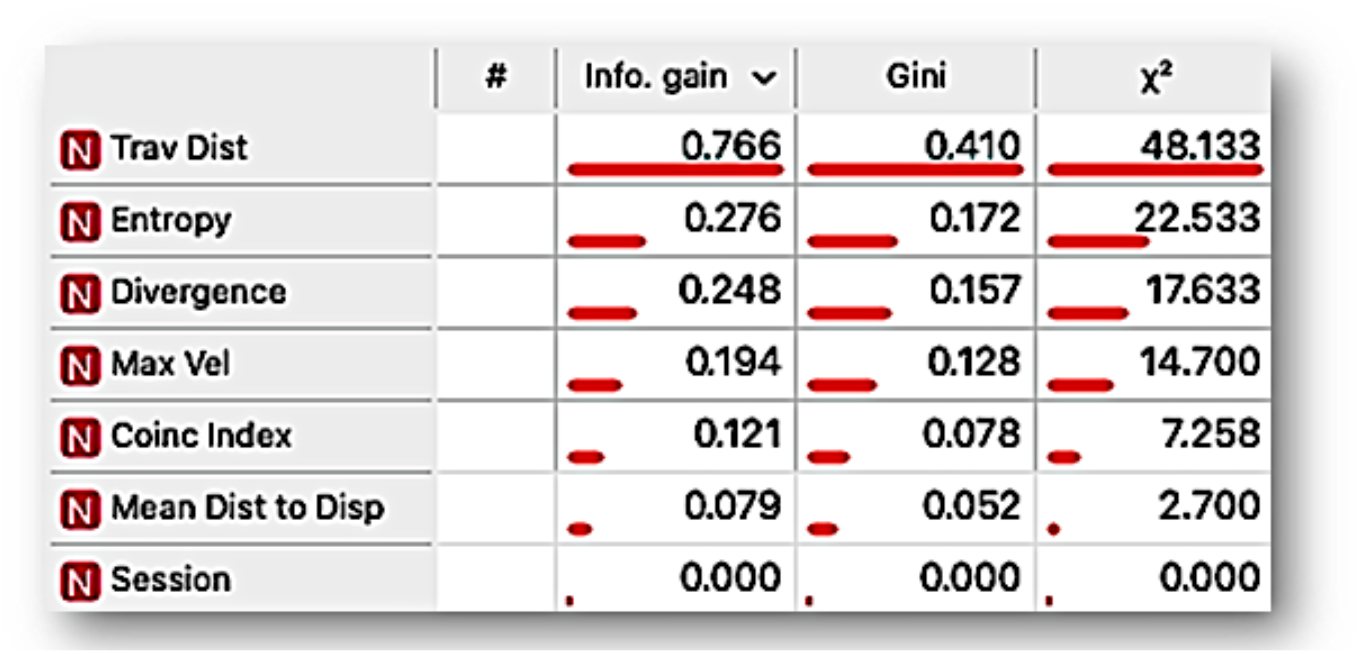
Ranking variable analysis, under *information gain, mean decrease impurity Gini index* and χ^2^ procedures, for the features: traveled distance per session (Trav Dist), entropy, divergence, maximum velocity per session (*Max Vel*), coincidence index (*Coinc index*), mean distance to the dispenser (*Mean Dist to Disp*), and session. The length of the bar ranks the features by the scoring value that measures the relevance of each feature to differentiate the behavioral system as a function of a given time based-schedule (Fixed vs. Variable Time).

To have a perspicuous representation that would allow identifying if the data, given its multiple dimensions, are articulated or grouped as a function of the kind of time-based schedule employed, we conducted t-distributed Stochastic Neighbor Embedding (t-SNE). t-SNE is a machine learning algorithm for the visualization of high-dimensional datasets into a bidimensional or three-dimensional space. t-SNE performs a non-linear dimensionality reduction task for embedding datasets and obtaining low dimension transformations as a result. Relations between high dimensional data, which might be impossible to observe due to a considerable amount of variables, could be distinguished after transforming them into a space with reduced dimension by t-SNE. Furthermore, the representation obtained by t-SNE is perspicuous because the data with similar values are closer to each other (in a low dimensional space, 2D or 3D) than data with dissimilar values (for a complete description of t-SNE, see supplementary material).

Figure 6 shows a representation, using t-SNE, for the data of all experimental sessions and subjects. Each point could be seen as multidimensional data for a session, considering the *distance traveled, entropy, divergence, maximum velocity, coincidence index, mean distance to the dispenser*, and *session number into the experiment* as dimensions, with the condition, FT vs. VT, as target feature. FT data (blue points) tended to be closer to each other, and the same was observed concerning VT data (red points). Color regions are shown in the figure to facilitate the visualization of groupings. The conformation of only two predominant and well-delimitated regions is clear, and FT and VT data are separated, except for a few dots inserted in the colored region of the opposite condition. Given that the multidimensional space of t-SNE could be seen as behavioral system representation, as a whole, the main finding is that emerged well-differentiated behavioral system under each condition or schedule.

**Figure 6.**
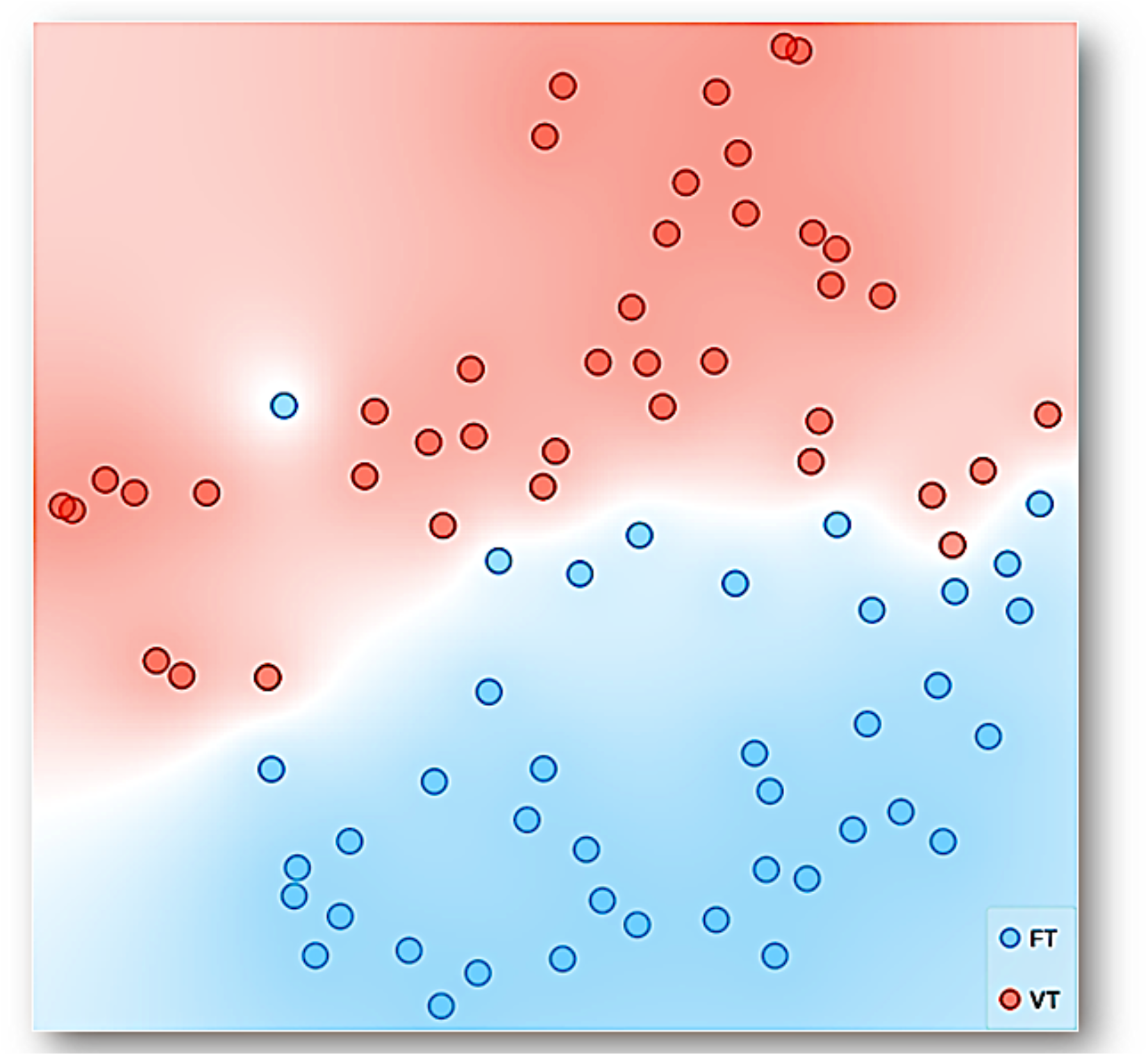
Representation with TSNE for the data of all experimental sessions and subjects. Each point represents multidimensional data for a session, considering: the *distance traveled, entropy, divergence, maximum velocity, coincidence index, mean distance to the dispenser*, and *session number into the experiment* as dimensions, with time based-schedule, Fixed Time (FT), and Variable Time (VT), as a target feature. The data with similar values, given the multiple features or input variables taken as a whole, is simply closer to each other than data with dissimilar values.

Figure 7 shows the linear projection (for a full description, see the supplementary material) of the multidimensional data of all sessions and subjects for six dimensions: *coincidence index, divergence, mean distance to the water dispenser, entropy, traveled distance*, and *maximum velocity*. The direction of each vector points out the direction to the increasing values for a given dimension. Colored regions related to each condition are added to facilitate the visualization of the data tendency. The prevalence of a colored region in a given dimension represents higher values for the correspondent experimental condition to such color compared to the other experimental condition. Thus, the linear projection showed higher *maximum velocity, traveled distance*, and *entropy* values under FT than VT. While showed values nearby for *mean distance to the dispenser* for both conditions, a red shadow suggests higher values for VT.

**Figure 7.**
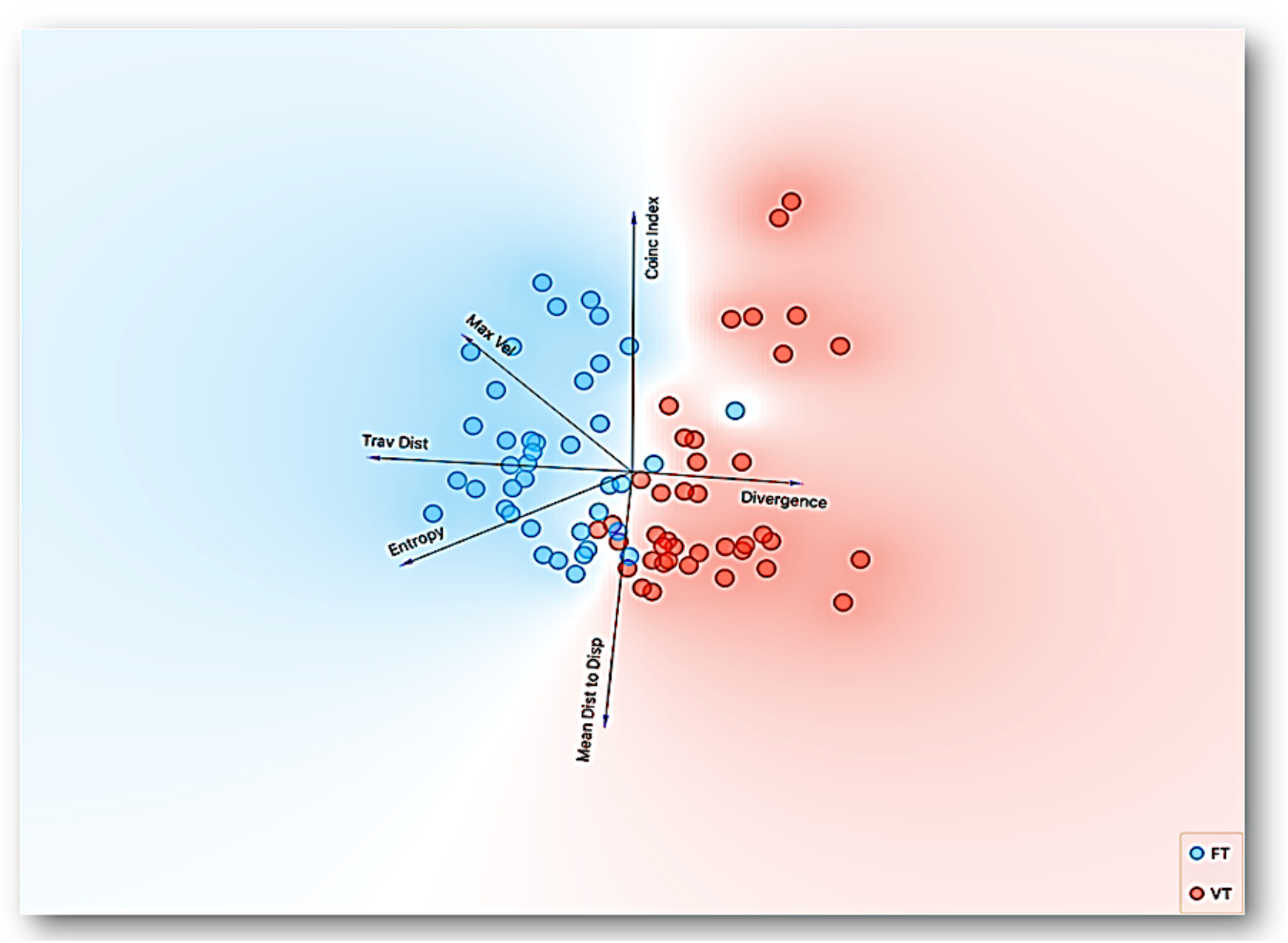
Representation with Linear Projection, by Principal Component Analysis, of multidimensional data of all sessions and subjects for six dimensions: *coincidence index, divergence, mean distance to the water dispenser, entropy, traveled distance*, and *maximum velocity*, with time based-schedule, Fixed Time (FT) and Variable Time (VT), as a target feature. The direction of each vector points out the direction to increasing values for a given dimension.

On the other hand, the projection clearly showed higher *divergence* values for VT and nearby values for the *coincidence index* for both conditions; nevertheless, the blue shadow in this last vector pointed out higher values for FT. One relevant difference of this representation with other reduction dimension procedures is that it has specific representations for each relevant dimension of the data in the orthogonal space. The linear projection representation confirms the relevance of the spatial dimensions of behavior related to the programmed time-based schedules.

### Example 2

#### Motivational operations: Behavioral dynamics under different deprivations in concurrent schedules

A procedure that is conducted in most studies of the experimental analysis of behavior that uses ‘appetitive’ stimulus is the deprivation of a given commodity (e.g., water or food) that it is later used to be delivered contingent to some response or behavioral pattern in contingent schedules (Skinner, 1938). In non-contingent schedules, the delivery of such commodity is presented in conjunction with given stimuli (e.g., pavlovian conditioning procedures) or simply presented according to a specific time rule (e.g., time-based schedules). The deprivation procedure, in methodological terms, has the purpose of establishing the relevance or dispositional value of the delivered commodity as stimulus (Michael, 1982; Reberg et al., 1978). This dispositional value is crucial to explaining behavioral systems, seen as articulating behavioral patterns, responses, and other stimuli. The study of the relevance or dispositional value of a given stimulus, *por mor* of the deprivation operation, has been related to the field of motivation, under the terms of ‘motivational operations’ (Laraway et al., 2003), ‘establishment operations’ (Michael, 1982, 1993), among others.

Several years ago, it was established that the ‘motivational’ function of a given stimulus could be characterized based on *direction, intensity*, and *variation* of the spatial behavior (Berlyne, 1955; De Valois, 1954; Duffy, 1951; 1957; Maier & Schneirla, 1964; Schneirla, 1959). Nonetheless, this characterization has been restricted only to the rate response of the discrete responses under the single response paradigm. Thus, the spatial dimension of the behavior and its dynamics has been ignored in the contemporary study of the EAB (e.g., Lewon et al. 2019).

On the other hand, generally, the effect of food or water deprivation is evaluated by removing access to them to the experimental subjects outside the experimental sessions and then presenting one or the other, either contingent or non-contingent, to a given response (Skinner, 1938; Bolles, 1975) during the session. The effect of presenting food and water concurrently when subjects are food or water-deprived has been less studied (Fallon et al., 1965, Lewon et al., 2019). Under this rationale, the objective of this study was to evaluate the effects of food and water deprivation conditions on the behavioral continuum in conditions where food and water are concurrently delivered. A multidimensional and multilevel analysis and data representation was conducted using machine learning tools to integrate standard discrete responses and spatial dynamics.

## Method

### Subjects

Five female and one male (Subject 3) Wistar rats (5 months old) were used. According to the current phase of the experiment, rats were housed in individual home cages and placed on a water or food-deprivation schedule for 22 hrs before every experimental session. All procedures were conducted according to university regulations of animal use and care and followed the official Mexican norm NOM-062-ZOO-1999 for Technical Specification for Production, Use, and Care of Laboratory Animals.

### Apparatus

An experimental chamber of 92 cm width × 92 cm long and 33 cm height was used. 2 cm above the grid floor and in the center of opposed walls, two dispensers were located, a liquid dipper (Coulbourn E14-05,) and a modified food receptacle with a pellet dispenser (Coulbourn E14-24). The dipper allowed access to 0.1cc of water for 3s, while the pellet dispenser delivered a 45 mg. pellet with limited availability of 3 s. Entries to both dispensers were detected by Head Entry Detectors (MED ENV-254-CB). In addition, above both dispensers, a yellow light was used as a visual stimulus (MED ENV-222M) to indicate food or water delivery (for a diagram of the apparatus, see Hernández, León, and Quintero, 2021).

Water and food deliveries were programmed and registered with the Software MED PC IV, and head entries were also registered using this software. Rat’s displacement in the experimental chamber was recorded using a video camera Topica TP-505D/3, 1m above the chamber. The video camera was connected to a PC with software Ethovision 2.3. With this software were obtained records of rats’ displacement in X, Y coordinates every 0.2 s.

### Procedure

#### Experimental phase

After an initial training phase to the food and water dispenser (see Hernández et al. for a complete description), subjects were exposed to two deprivation conditions: 1) Water-Deprivation (WD) and 2) Food-Deprivation (FD). Each deprivation consisted of three days with the corresponding food or water restriction and one experimental session per day. After each condition, subjects were allowed unrestricted access to both commodities for 24 h before the following deprivation condition to avoid a drastic decrease in weight and separate the effect of each deprivation (Lewon et al., 2019). Subjects were assigned to one of two deprivation sequences to control for the potential effect of the first deprivation condition on the following condition. The specific order of deprivations for each sequence is shown in Table 1. All experimental sessions consisted of presenting a CONC FT 30s FT 30s schedule of food and water with limited availability of 3s. A yellow light above both dispensers was turned on with every delivery and remained during food or water availability. All experimental sessions lasted 30 min.

**Table 1.** Sequence of deprivation conditions for each group.

| Sequence | Deprivation Condition |  |
| --- | --- | --- |
| 1 | WD | FD |
| 2 | FD | WD |

*Note.* WD: Water Deprivation, FD: Food Deprivation. Each condition lasted three sessions, and each session lasted 30 minutes. Three rats were assigned to Sequence 1 and three rats to Sequence 2.

### Data analysis

The same analytical approach of Experiment 1 was used in Experiment 2, only with appropriate settings due to the differences in methods, apparatus, and records. Specifically, measures related to water and food dispensers were added, such as entrances and derivated measures (*intensity, precision to dispensers*, and *proportion* to commodities contacted). Each measure is described in the following section.

## Results

We conducted a multilevel analysis to characterize the behavioral continuum and compare the spatial dynamics of behavior under water and food deprivation. In the same way of Example 1, first, we show representative within-subject results of the behavioral continuum within-session, for the initial, intermediate, and last session of the experiment, for one representative subject. Second, we show summary results for measures based on spatial behavior, first and non-first order, and measures based on discrete response in water and food dispensers. Third, we conducted a multidimensional analysis and integrative representations based on Machine Learning.

Figure 8 shows the continuum spatial-behavioral data for the complete initial, intermediate and final sessions for a representative experimental subject. Panel A represents the routes of the subject. There was a higher variation of spatial behavior at the arena under Water Deprivation (WD) than under Food Deprivation (FD) for the intermediate and final sessions. In addition, higher spatial behavior in the food dispenser zone (top of the plot) under FD and a distributed densification between both dispenser zones (top and down of the plot) under WD were observed. Panel B shows the distance to the food dispenser (red line) and water dispenser (blue line), moment to moment (each .02 sec). For the intermediate and final sessions under FD, a small distance to the food dispenser and a long distance to the water dispenser was observed, with only a few alternations between high and low distance values to both dispensers.

**Figure 8.**
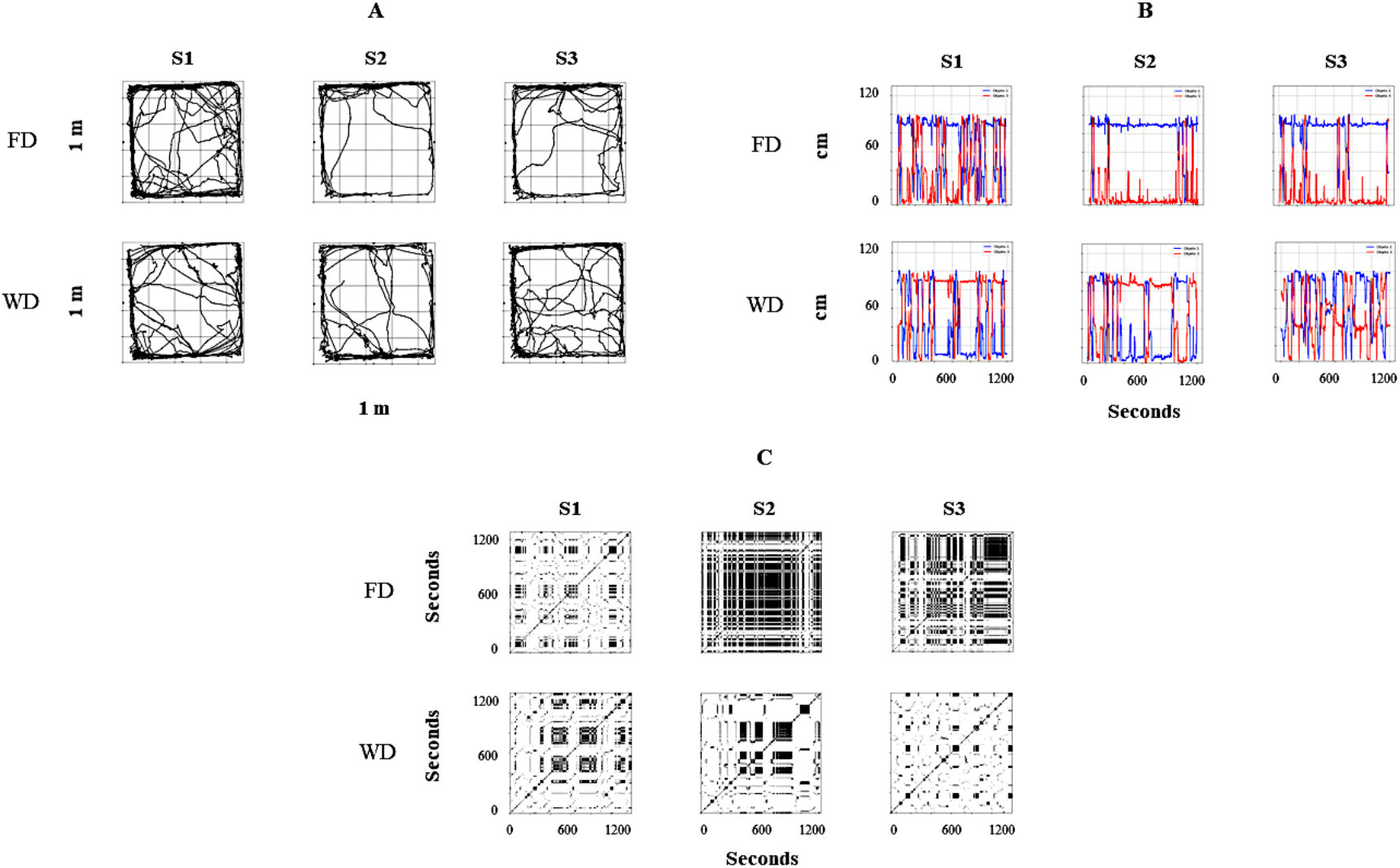
Representative within-subject results for all sessions of the behavioral continuum for one rat (R5). Each column depicts a session, and rows depict deprivation conditions (FD and WD, respectively). Panel A shows the routes of the rat in a bidimensional representation of the experimental chamber; red circles represent Food Dispenser location, blue circles represent Water Dispenser location. Panel B depicts the distance to the dispensers (Y-axis) moment-to-moment (X-axis); the red line represents the distance to Food Dispenser; the blue line represents the distance to Water Dispenser. Finally, panel C shows recurrence plots; this changes regions of each rat (in a matrixial configuration of 10 × 10 virtual zones) as the session progresses.

In contrast, under WD, for the intermediate and final sessions, a significant alternation between high and low values of distance to the dispensers and, then, a clear back and forth pattern between dispensers was observed. Panel C depicts the recurrence plots (see Figure 2 description and supplementary material). High permanence (extended black zones), with only some transitions, under FD and high recurrence (black-white mosaic patterns) under WD were observed in these plots. Thus, the three panels (A, B & C), as a whole, suggest a robust difference in the spatial dynamics by Food Deprivation vs. Water Deprivation, under the same concurrent schedule and for the same experimental subject.

Figure 9 shows summary results for the measures based on the spatial dimension of behavior: *mean distance to the food dispenser, mean distance to the water dispenser, mean distance to the center of the experimental arena, entropy index, and divergence index*, under Food Deprivation (FD) and Water Deprivation (WD), for all sessions and experimental subjects, independently of the sequence in which they were exposed. All measures were sensitive to the deprivation condition, except the divergence index. The distance to the food dispenser had low values under FD and relatively high values under WD. In contrast, the opposite effect concerning the water dispenser was observed, with relatively low distance values under WD and high values under FD. The data of distance to the dispensers was more spread under WD than under FD. The mean distance to the center was higher under WD than FD, and the entropy index too. All the previously mentioned findings were robust and point out a significant differential spatial dynamic of the behavior related to deprivation conditions.

**Figure 9.**
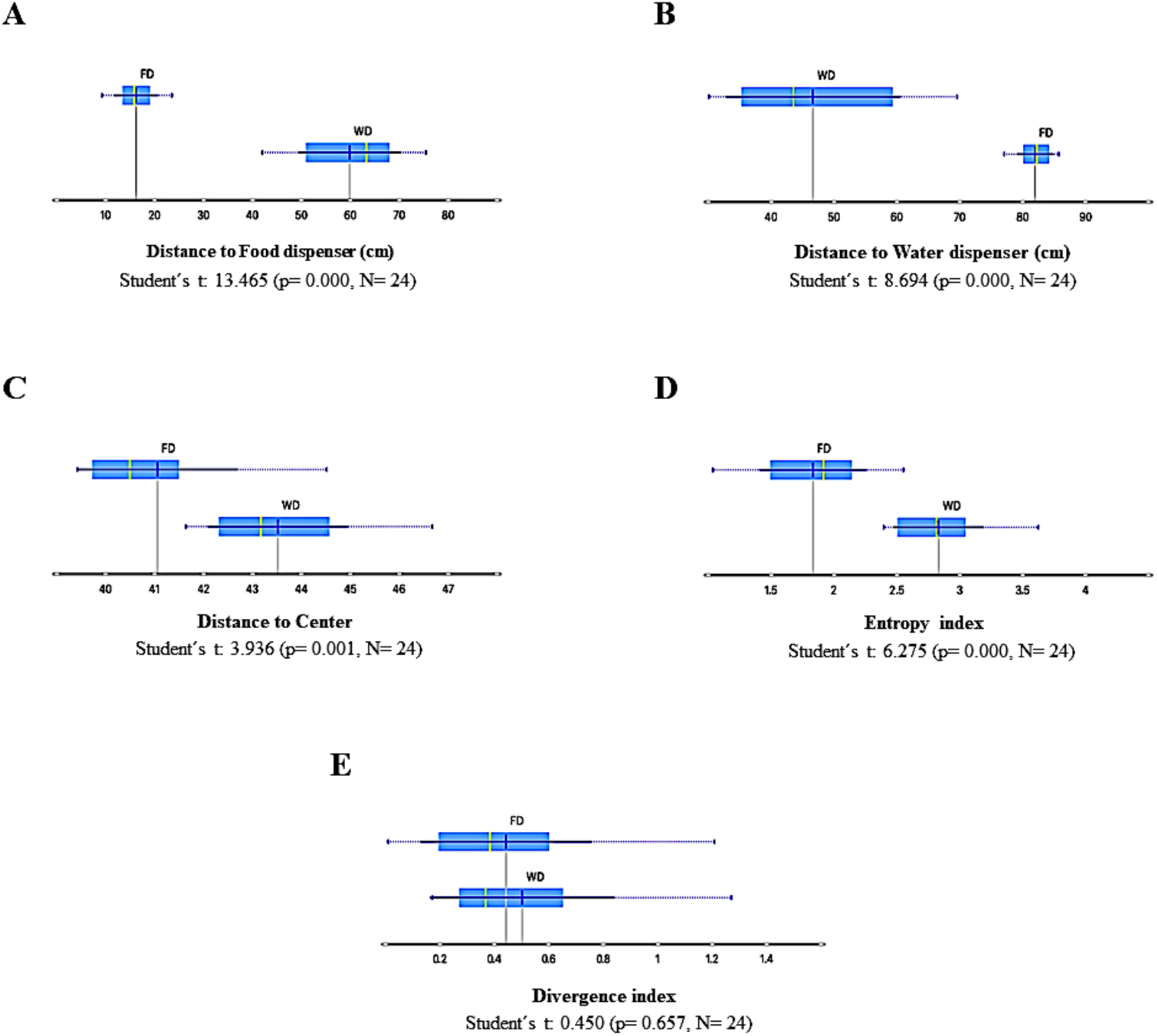
Summary results of spatial-behavior measures for all sessions and subjects under Food Deprivation (FD) and Water Deprivation (WD) schedules. Panel A, Distance to food dispenser; Panel B, Distance to water dispenser; Panel C, Distance to center; Panel D, Entropy; Panel E, Divergence. Each box depicts the mean (dark blue vertical line), the median (yellow vertical line), the standard deviation (thin blue line), and the values between the first and the third quartile (blue highlighted area).

Figure 10 shows a summary results for measures based on discrete responses, namely *a) intensity to food dispenser index*, b) *intensity to water dispenser index*, c) *precision to food dispenser index*, d) *precision to water dispenser index*, e) *proportion to food contacted*, and f) *proportion of water contacted*. These measures are relevant because they are related to those employed in the standard paradigms based on single discrete responses. Panel A and Panel B depict the intensity related to the dispensers; the *intensity index* was obtained by dividing the total number of head entries in one dispenser of each session by the maximum number of entries in any session on the whole experiment for a particular subject and dispenser. Then this procedure was carried on for each subject and session, always using this within-subject and within-dispenser comparison. The *intensity to the food dispenser* was high under FD and low under WD, while the *intensity to the water dispenser* was high, though spread, under WD and very low under FD. Panel C and Panel D show the precision to the dispensers. *Precision index* was obtained by dividing the number of entries to each dispenser, when the commodity was available (water or food), by the total number of head entries in that session and in that dispenser. The data below percentile 5 for each subject on each dispenser were eliminated to palliate *possible ceiling* or *floor effects* in sessions. The *precision related to the food dispenser* was lower under FD than under WD, and there was no difference in precision to water dispenser between deprivations (FD and WD). Finally, Panel E and Panel F depict the proportion to food and water contacted from the total available. The *proportion to food contacted* was higher under FD than WD, while *water contacted* was lower under FD than WD. The measures based on discrete responses, as a whole, show interesting findings. First, the *intensity* and *effectiveness* (i.e., proportion to food contacted) to each dispenser clearly depend on the deprivation condition; this is, high values of intensity and effectiveness to food dispenser was observed under FD, and vice versa, high values of intensity and effectiveness to water dispenser was observed under WD.

**Figure 10.**
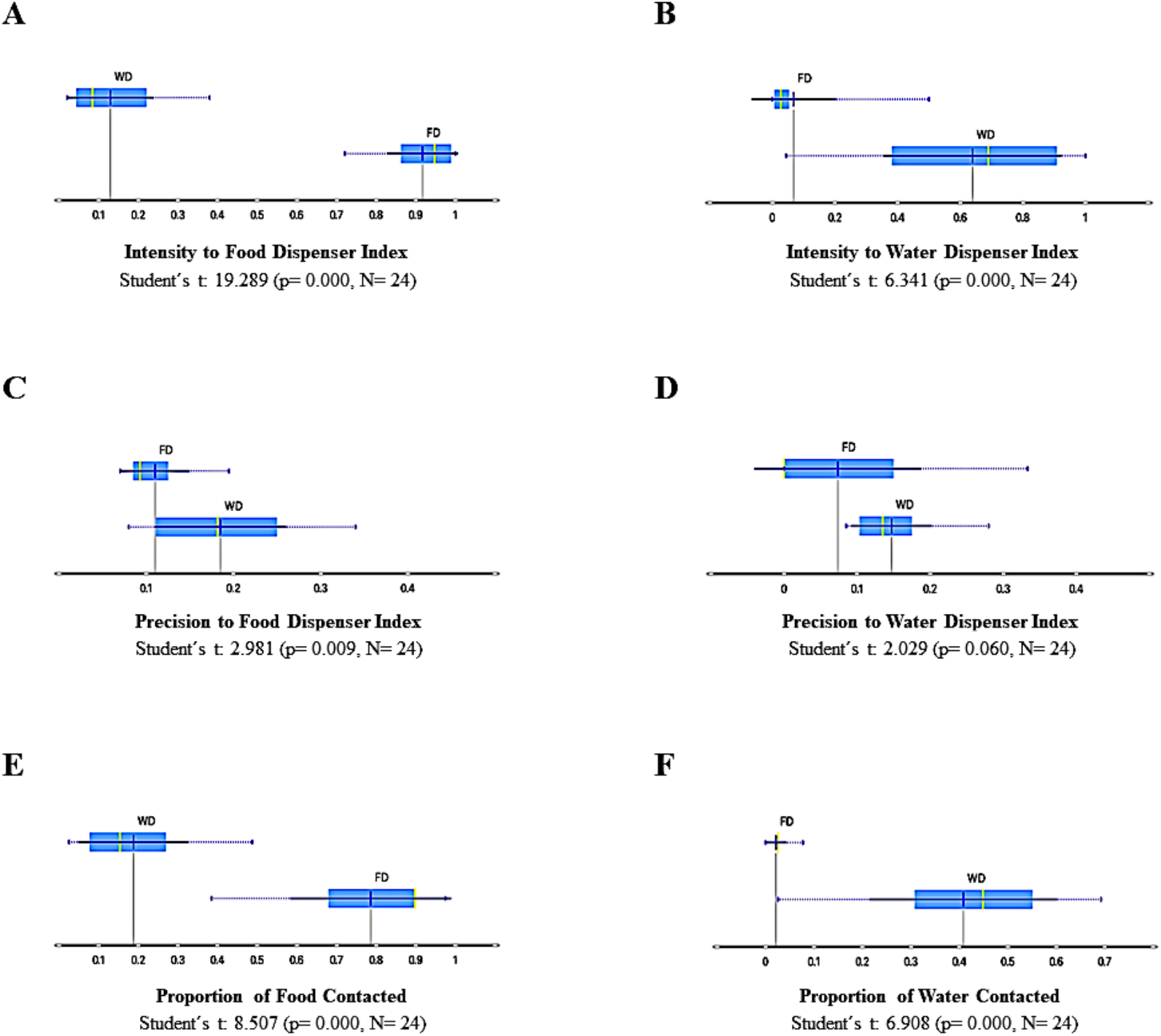
Summary results of measures based on discrete responses, for all sessions and subjects under Food Deprivation (FD) and Water Deprivation (WD) schedules: Intensity to food dispenser index (Panel A), intensity to water dispenser index (Panel B), Precision to food dispenser index (Panel C), Precision to water dispenser index (Panel D), Proportion of food contacted (Panel E) and Proportion of water contacted (Panel F). Each box depicts the mean (dark blue vertical line), the median (yellow vertical line), the standard deviation

Nevertheless, the *modulating effect* of each deprivation condition over the behavior related to the correspondent dispenser and commodity delivery to such deprivation is not precisely the same for both deprivations. On the one hand, the *modulating effect* of FD, for all behavioral measures based on discrete responses, is most robust for the food dispenser than the effect of WD over the same measures related to the water dispenser. On the other hand, the data under FD tends to extreme values (very low or very high) related to *intensity* and *effectiveness*, while under WD tends to intermediate and spread values. Finally, it is remarkable that the precision of behavior related to food delivery was negatively affected under FD. This effect for water delivery under WD was not observed.

Figure 11 shows the variable-ranking under three procedures, namely *Information gain, impurity Gini index* and χ^2^, for all measures, those based on continuum spatial dimension of behavior and those based on discrete responses (for a complete description concerning these algorithms, see the supplementary material, and for suggested use in behavior analysis, see the description of Figure 5). The variable ranking allows identifying the relevance of each variable (i.e., behavioral measure or dimension) into the whole multidimensional system. Given our subject matter, the modulating effect of two different deprivations (FD and WD) into the multidimensional behavioral system, the variable ranking suggests that the most relevant variables were the distance to the dispensers, intensity to the dispensers, and entropy. These findings suggest that the spatial dimension of behavior was as relevant as discrete responses into the behavioral systems that emerge by modulation under food deprivation and water deprivation.

**Figure 11.**
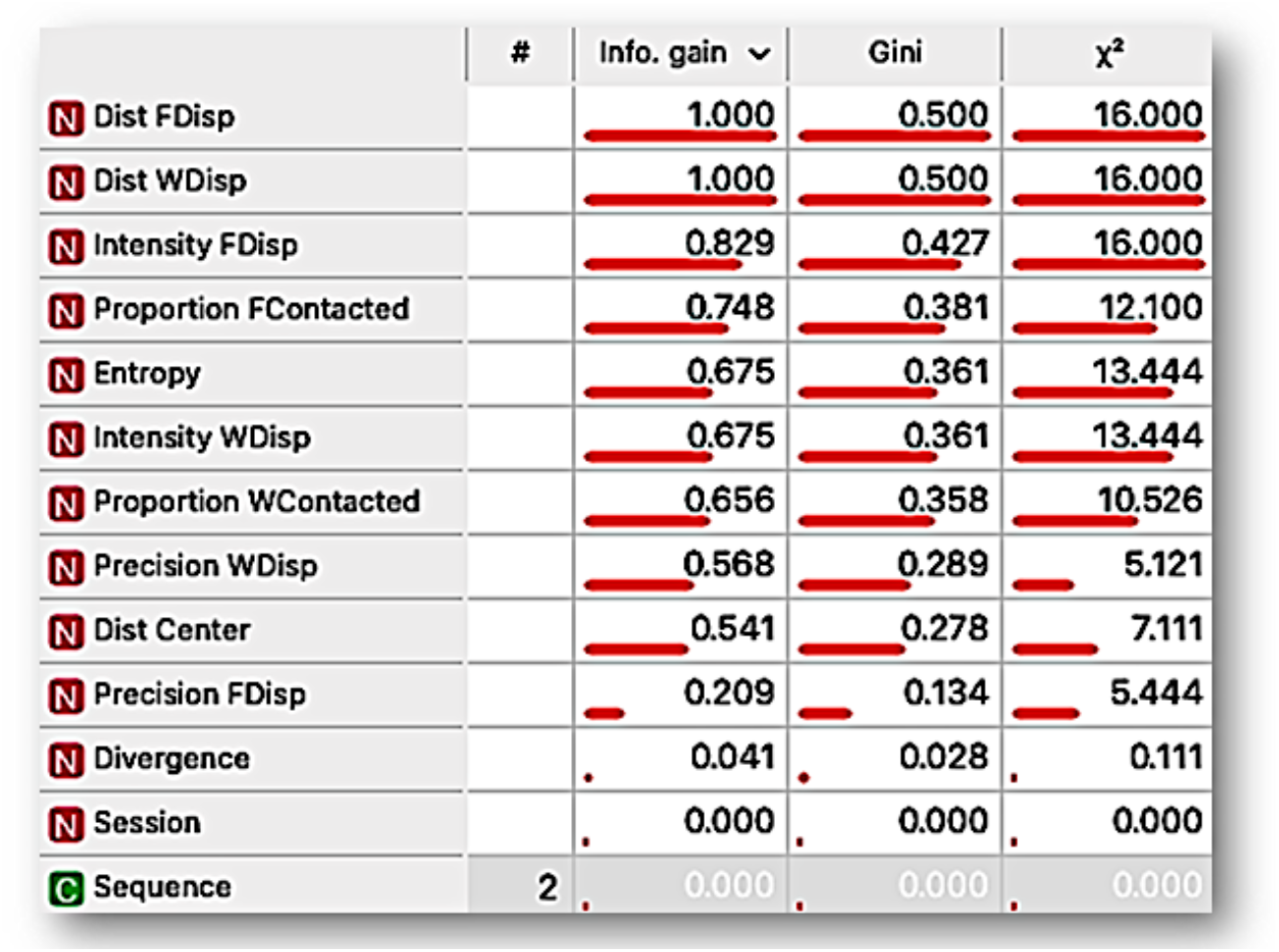
Ranking variable analysis, under *information gain, mean decrease impurity Gini index* and χ^2^ procedures, for the features: Distance to food dispenser (*Dist FDisp*), distance to water dispenser (*Dist WDisp*), intensity to food dispenser (*Intensity FDispenser*), Proportion of food contacted (*Proportion FContacted*), Entropy, Intensity to water dispenser (*Intensity WDisp*), Proportion of water contacted (*Proportion WContacted*), Precision to water dispenser (Precision WDisp), distance to the center of the experimental arena (*Dist Center*), Precision to food dispenser (*Precision FDisp*), Divergence, Session, Sequence.

Figure 12 shows a representation by t-SNE (for a complete explanation, see supplementary material and description of Figure 6) for the data of all the experimental sessions and subjects. Each point could be seen as multidimensional data for a session, considering both measures based on spatial behavior continuum and discrete responses (see Figure 11) and deprivation condition (FD and WD) as the target feature. Data tends to be closer by condition for both FD (red dots) and WD (blue dots). Two well-delimitated colored regions were formed, with a clear separation between FD and WD data. Additionally, two clusters were conformed under the K-means clustering procedure (for a description, see supplementary material), see Panel A. The coincidence of each cluster data with a deprivation condition data was very robust. This analysis could be taken as an explicit confirmation of the differential modulation by each deprivation condition (FD vs. WD) over the emerged multidimensional behavioral system under the same concurrent schedule and within-subject design. Finally, in Panel B, another complementary representation was conducted. In this panel, the circles point out the deprivation sequence 1 (WD-FD), and the crosses sign out the deprivations sequence 2 (FD-WD). The representation shows that the data clustering by deprivation was robust to the sequences (that is, there are well-delimited regions for each deprivation regardless of the sequence), but also suggests that the data tended to be close by sequence within deprivation. In addition, the representation suggests a *contrast effect* on deprivation conditions by Sequence 1; this is, the FD data and WD data were more distant from each other than the data for the same deprivation conditions in Sequence 2.

**Figure 12.**
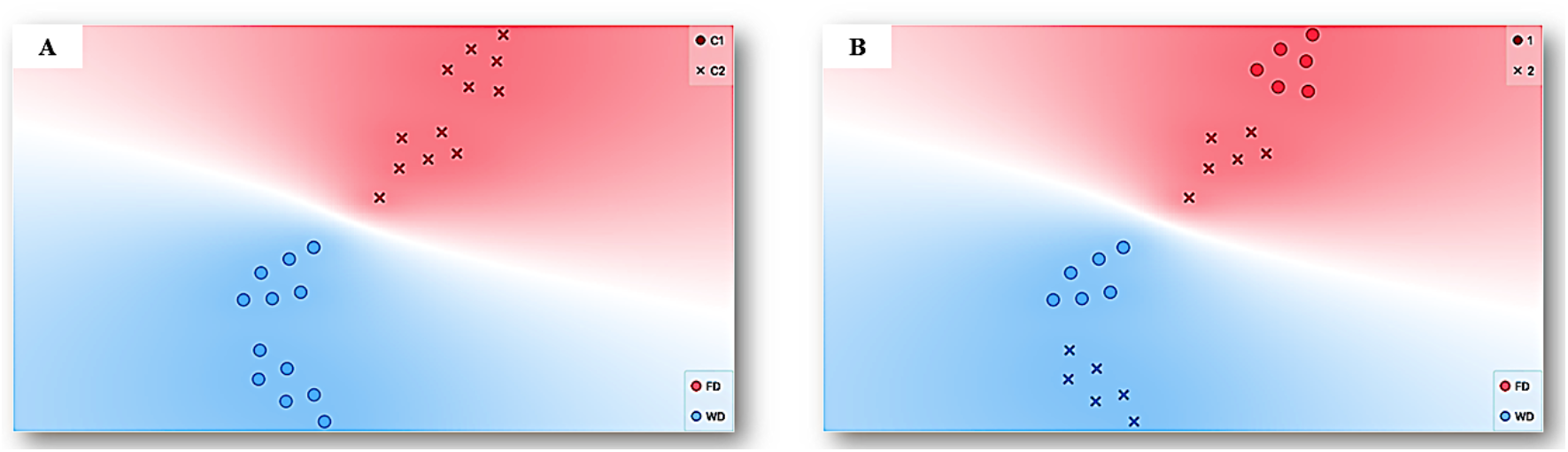
Representation with TSNE for the data of all experimental sessions and subjects. Each point represents multidimensional data for a session, given all features (see Figure 11), with deprivation condition, Food Deprivation (FD-red), and Water Deprivation (WD-blue), as a target feature. The data with similar values, given the multiple features or input variables taken as a whole, is simply closer to each other than data with dissimilar values. Panel A, additionally, shows clustering under the K-means procedure and two well-delimitated colored regions, with a clear separation between FD and WD data, and corresponding each one with a cluster (C1 -circles- and C2 -crosses-). On the other hand, Panel B depicts the deprivation sequence (Sequence1 -Circle- and Sequence2 - crosses-).

## Discussion

The purpose of the present work was threefold: 1) to propose an integrative and multidimensional approach for the analysis of behavioral systems; 2) to show novel behavioral aspects revealed under a multidimensional approach based on the integration of discrete and continuous data assisted by Machine Learning tools; and 3) to provide relevant and novel behavioral measures and data representations based on the integration of spatial dynamics and discrete responses, for the study of behavioral systems related to relevant research areas in behavioral science such as *water-seeking behavior* and *motivational operations*.

In the first example, concerning behavioral dynamics under FT and VT, marked differences in *routes, rat’s location at the moment of water delivery, distance to the dispenser*, back and forth to the dispenser, and *recurrence patterns* were observed. These findings suggest a considerable difference in emergent spatial behavior (*direction* and *variation)* under both temporal schedules (Fixed vs. Variable Time). In addition, they confirm our hypothesis that the proposed first-order measures based on spatial behavior will be sensitive to EAB paradigmatic procedures.

Furthermore, the *entropy*, a non-first order measure, was sensitive to the programmed contingencies, with higher values under FT than VT; the behavioral meaning of this finding is that the distribution of the organism location presents more variability under FT than VT. This finding is interesting because the temporal variation in water delivery was associated with lower variability of organism location, and temporal constancy or fixation was associated with higher variation of the organism location. On the other hand, the organism’s location variability distribution showed a low *divergence* between sessions under FT and not under VT. As far as we know, the use of *entropy* and *divergence* to characterize the spatial variability of behavior is scarce. Nevertheless, our findings revealed that *entropy* and *divergence* are embedded features of spatial behavior with a higher sensitivity to the temporal schedules.

The findings together, of first and non-first-order measures, with data of *direction* of behavior, such as routes, back and forth patterns; and others about the *variation* of behavior, such as checker recurrence patterns, and higher values of entropy under FT, could be seen as an objective measure of the idiosyncratic spatial patterns reported anecdotally in the literature as superstition behavior under FT (Skinner, 1948). Thus, in our perspective, the analysis carried out and its behavioral meaning shows the plausibility and parsimony of the CEAB approach.

On the other hand, the analysis of the different features and its ranking variable assisted by Machine Learning confirmed our hypothesis, related to the relevance of spatial features over standard discrete responses (e.g., water contacted or coincidence index) in the behavioral systems under temporal schedules. The ranking variable analysis shows that traveled distance, entropy, divergence, and maximum velocity were more sensitive to the programmed schedules than the standard feature of water contacted, measured as a coincidence index. Finally, t-SNE and Linear Projection were helpful to represent multidimensional-behavioral systems in a perspicuous way (e.g., in bi-dimensional space). These representations allow confirmation that each schedule (FT and VT) gives place to well-differentiated behavioral systems based on spatial behavior and a discrete response.

In the second example, concerning behavioral dynamics in concurrent schedules under different deprivation conditions (WD vs. FD), more extended routes, back and forth patterns alternated between dispensers, and recurrence patterns were observed under Water-Deprivation than under Food Deprivation. Again, findings suggest these representations were sensitive to the deprivation condition.

Furthermore, all measures related to spatial behavior were markedly affected by deprivation conditions (e.g., *distance to both dispensers*; *distance to the center of the arena* and *entropy*), except *divergence*. The latter indicates consistency between sessions related to the variability values of the organism’s location distribution under both deprivations. As in Example 1, the findings show that *entropy* is a relevant feature embedded in spatial behavior that is significantly affected by a standard procedure with well-known effects. In simple words, the findings suggest spatial behavior is very sensitive to namely ‘motivational operations’ under choice situations (e.g., concurrent schedules). A relevant aspect is that these features are indicators of *direction* and *variability* of the behavior that could be used as an alternative indicator to identify the motivational function of a given procedure aside from discrete responses.

On the other hand, all measures based on discrete responses were sensitive to deprivation conditions, except the precision to the water dispenser. In general terms, each deprivation condition affected the *direction* of behavior, both spatial and discrete responses, to correspondent commodity. Although, the effect was not exactly symmetrical. These findings are consistent with the expected under the standard paradigm and the literature. This point is crucial because it increases the validity of our findings and conclusions concerning spatial behavior (as a simile of concurrent validity in a non-statistical way).

The ranking variable analysis, assisted by Machine Learning, considered eleven features (five based on spatial behavior and six on discrete responses). It reveals that three are related to the spatial behavior of the five most relevant features: *distance to the food dispenser, distance to the water dispenser*, and *entropy*. These findings confirm our hypothesis related to the CEAB will reveal that spatial features are at least as relevant as behavioral features based on discrete responses, but now concerning other behavioral phenomena and paradigms, ‘motivation’ and ‘motivational operations under concurrent schedules’ respectively. Finally, t-SNE showed that each deprivation condition gives well-differentiated behavioral systems based on spatial behavior and a discrete response under the same concurrent schedules.

As our examples show, the general proposed approach in this work helped integrate a multiple-level analysis to coalesce discrete and continuous dimensions of behavior (and derivate first and non-first order measures) as a whole system. It also proved fruitful to provide a broad characterization of the continuum of behavior in which the spatial dynamics are on the first plane. The proposed approach appears promising to characterize and integrate different behavioral features as a whole behavioral system, pointed to as relevant throughout the development of behavioral science. Among these features are *direction* (Schneirla, 1959; in our work distance to the dispenser), *intensity* (Duffy, 1957; in our work speed, acceleration), *variation* (Antonitis, 1951; Berlyne, 1955; Iversen, 2017; Mowrer & Jones, 1943; in our work entropy), *preference* (Irwin, 1958; in our work time spent in a given zone), *persistence* (Bolles, 1975; in our work dispenser entries). So, with the proposed multidisciplinary methodological approach, the purpose of overcoming the segmented characterization of the behavioral continuum and its derived paradigms, for example, the single response paradigm (Henton & Iversen, 1978), could go beyond the theoretical level that has been maintained up to now (Kantor, 1958).

The findings of both experiments, presented to exemplify our approach, show that the recording and analysis of the continuum of spatial-behaviour of the organisms is of primary importance to account for the principles that underlie in behavioural systems, and suggest that: a) moment-to-moment analysis and representations of locomotion-based data, across complete sessions, are helpful to identify, and characterize, the behavioral dynamics under different stimuli-schedules and deprivation conditions (see *routes, distance to the dispensers* and *recurrence plots*); b) the proposed non-first order variables (i.e. *entropy* and *divergence*), based on locomotion data, are relevant and sensible to stimuli-schedule and deprivation conditions; c) the variables based on locomotion-data could be more sensible, than variables based on discrete responses, to stimuli schedules and deprivation conditions (see *variable-ranking* analysis based on Machine Learning); d) discrete responses and the continuum of spatial behaviour comprise an unitary and whole system, that could be apprehend and represent, in a perspicuous way, with Machine Learning tools like *t-SNE, clustering* based on *K-means, linear projection*, among others.

Our examples and findings suggest that the proposed multidisciplinary approach (CEAB) allows going forward on explaining behavioral systems and reveals an integration of spatial dynamics and discrete responses hidden until now for the behavioral science. In addition, new empirical relations and insights were revealed under the CEAB related to water-seeking behavior (Leon et al., 2020a) and motivational operations (Hernandez et al., 2021; Michael, 1982, 1993).

Although the proposed approach appears to be promising, to confirm its heuristic and parsimonious value, it should be evaluated under a) other relevant phenomena; b) other kinds of schedules (e.g., contingent schedules); c) different stimulating conditions (e.g., aversive stimulation); d) different organization of behavior (e.g., behavior under stimulus control; relational behavior); e) different species, including humans.

Finally, the proposed approach could be strengthened by integrating additional first and non-first-order measures pertinent to apprehend and characterize the dynamics of relevant dimensions of behavior (such as direction, variation, vigor, among others). On the other hand, additional Artificial Intelligence tools, like predictive analysis, could be explored to extend the scope of our approach for behavioral science, specifically for the experimental analysis of behavior.

As a corollary, the fast-paced development of contemporary computational tools of fields as Artificial Intelligence has rapidly changed the landscape of some fields of behavioral science in the last decades, for example, ethology (Dell et al., 2014) and neuroscience (Datta et al., 2019; Mathis et al., 2019; Mathis & Mathis, 2020; Wiltschko et al., 2015). It is time the non-mediational, systematic, parametric (Henton & Iversen, 1975; Schoenfeld & Cole, 1972; Skinner, 1938) and ecological (Silva & Timberlake, 1997; Timberlake, 1994) approaches in the experimental analysis of behavior start to profit from these tools (Turgeon & Lanovaz, 2020).

