## Supplementary material for "Beyond single discrete responses: An integrative and multidimensional analysis of behavioral dynamics assisted by Machine Learning": Suplementary Material (from another version of this preprint: version 1, 2021-03-17)

**Smoothed distance**

If $\left\{ d_{i} \right\}$ is the sequence of the distance function, where the index $i$ takes values from 1 to the maximum of data N, then the values for the smoothed distance function sequence $\left\{ \bar{d}_{i} \right\}$, are determined as follows, for a radius of K frames around each value. For $K<i<N-K$,

$$\bar{d}_{i}=\frac{1}{2K+1}\sum_{j=i-K}^{i+K} d_{j}.$$

The first K terms of $\{\bar{d}_{i}\}$, for $1\leq i\leq K$, are determined by

$$\bar{d}_{i}=\frac{1}{K+i}\sum_{j=i}^{i+K} d_{j}.$$

While the last K terms of $\{\bar{d}_{i}\}$, for $N-K\leq i\leq N$, are defined by the formula

$$\bar{d}_{i}=\frac{1}{K+N-i}\sum_{j=i-K}^{N} d_{j}.$$

**Recurrence**

The recurrence plot is defined by symmetric matrix $A=\left[ a_{ij} \right]$ with dimensions $N\times N$, in which the inputs are determined by the following function:

$$a_{ij}=\left\{ \begin{matrix} \text{black}, & \text{if }P_{j}\in R_{k} \text{and }P_{i}\in R_{k}\text{, } \\ \text{white}, & \text{if }P_{j}\notin R_{k} \text{and }P_{i}\in R_{k}, \end{matrix} \right.$$

where $i=1,2,\ldots,N$; $j=1,2,\ldots,N$; $k=1,2,\ldots,100$. That is, the matrix $A$ (recurrence plot) is a matrix of time (frame=.2 sec) per time (frame=.2 sec), from i to j. $P_{j}$ is a given frame and $R_{k}$ is a given region (one of hundred regions). If given frames $P_{i}$ & $P_{j}$ coincide in $R_{k}$, then the value $a_{ij}=$black in the intersection between $P_{i}$ & $P_{j}$ in the matrix $A$. If given frames $P_{i}$ & $P_{j}$ do not coincide in $R_{k}$, then the value $a_{ij}=$white in the intersection between $P_{i}$ & $P_{j}$ in the matrix $A$.

**Entropy**

The concept of entropy was introduced by Shannon (1948). Entropy is a measure associated with a discrete random variable, which indicates ‘uncertainty’ or ‘disorder’. The entropy measures the uncertainty average level of a random variable. When the variable has a low-probability value, the event carries more ‘information’ than when the variable produces a high-probability value. This notion of ‘information’ is also sometimes interpreted as “surprisal".

Formally, given a discrete random variable $X$ with possible outcomes $\left\{ x_{i} \right\}$, each with probability $P\left( x_{i} \right)$, the entropy $H\left( X, P \right)$ is as follows

$$H\left( X,P \right)=-\sum P\left( x_{i} \right)ln\left( P\left( x_{i} \right) \right).$$

It can be proven that the entropy of a discrete random variable is a non-negative number, $H\left( X,P \right)\geq0$, and its measure should be maximal if all the outcomes are equally likely (uncertainty is highest when all possible events are equiprobable).

In order to analyze the displacement pattern of individuals in each session, the discrete random variables $\left\{ x_{i} \right\}$ are the permanence in each defined zone from a configuration of $10\times10$ defined zones and $P\left( x_{i} \right)$ is accumulated time (standardized) at it.

**Divergence**

The Kullback-Leibler divergence was introduced by Kullback & Leibler (1951) and discussed by Kullback (1959). The Kullback-Leibler divergence (or relative entropy) is a measure of difference from first one probability distribution to second one.

For discrete probability distributions $P$ and $Q$ defined on the same random variable $X$, with possible outcomes $\left\{ x_{i} \right\}$, the Kullback–Leibler divergence from $Q$ to $P$ is defined as

$$D_{KL}\left( P\parallel Q \right)=\sum_{i} P\left( x_{i} \right)ln\left( \frac{P\left( x_{i} \right)}{Q\left( x_{i} \right)} \right).$$

The Kullback-Leibler divergence is defined only if for all $x$, $Q\left( x \right)=0$ implies $P\left( x \right)=0$. Whenever $P\left( x \right)$ is zero the contribution of the corresponding term is interpreted as zero.

The KL divergence $D_{KL}\left( P\parallel Q \right)$ can be thought of as something like a measurement of how far the distribution $Q$ is from the distribution $P$, because it is always non-negative ($D_{KL}\left( P\parallel Q \right)\geq0$) and a result known is $D_{KL}\left( P\parallel Q \right)$ zero if and only if $P=Q$, this is a Kullback–Leibler divergence of $0$ indicates that the two distributions in question are identical. However, it is not symmetric, that is, $D_{KL}\left( P\parallel Q \right)\neq D_{KL}\left( Q\parallel P \right)$.

To analyze the displacement pattern of individuals in consecutive sessions, the discrete random variables $\left\{ x_{i} \right\}$ are the permanence in each square region from a configuration of $10\times10$ defined zones and $Q\left( x_{i} \right)$ is the accumulated time (standardized) at the first session and $P\left( x_{i} \right)$ is the accumulated time (standardized) at second one.

**Variable ranking**

The variable ranking is a lenient version of feature selection which consists of a process of ordering a set of features (input variables) normally by the value of a scoring function that measures the relevance of each feature according to a target (or output variable) as a predicting tool. Feature selection could be too rigid for some applications where determining the variables that must be removed or preserved is not clear given that all features are required to explain or predict a target value, but the relevance of some characteristics is more important for the purposes of the investigation. Even in some cases, variable ranking is the first phase for variable selection.

The algorithms for variable ranking are divided into two main classes: wrappers and filters. On the one hand, we can find several wrappers algorithms, most of them rely on training a machine learning algorithm like SVM or Decision Tree to estimate the quality of a subset of features of the dataset in order to predict the target variable; this approach could consume a large amount of computing resources for high-dimensional data sets, when looking for a minimal optimal feature set.

On the other hand, filter algorithms are the most widely used, due to the low computing resources used to apply them, even on high dimensional datasets. There are versions for single or multiple variable classification and their implementation also depends on whether the target variable is numerical or categorical. Since, in our experiments, the data sets have only categorical targets, we opted for this approach. Three theoretical information filter algorithms, conducted using Orange© 3.26, were used for the classification of a single variable ranking: information gain, mean decrease Gini, and $\boldsymbol{\chi}^{\mathbf{2}}$.

We will denote by T the set of training samples in the form of tuples $\left( x,y \right)=\left( x_{1},x_{2}, x_{3},\ldots, x_{n},y \right)$, where $x_{a}$ is the value of the $a^{th}$ feature (e.g., mean distance to relevant object, mean velocity, entropy and so on) of the example x and y is the corresponding target variable to a class label (e.g. water deprivation; food deprivation).

**t-SNE**

The t-distributed Stochastic Neighbor Embedding is a Machine Learning algorithm commonly used for visualization of high-dimensional datasets into a 2D o 3D space. In broad terms, t-SNE performs a non-linear dimensionality reduction task for embedding datasets and obtaining low dimension transformations as result, suitable for visualization where data entries with similar values for the features taken as inputs would be closer to each other than entries with dissimilar values, relations between inputs that might be impossible to observe due to the large number of variables would be distinguished after transforming them into a space with reduced dimension. The t-SNE analysis were conducted using Orange© 3.26

The t-SNE algorithm performs two phases to transform input data, first it builds a probability distribution with each pair of high-dimension data entries looking for similar data to obtain a higher probability and placing them closer, and a lower probability for data with values than have a significant difference between them. The second phase consists in defining a probability distribution over a reduced dimension space (2D or 3D) and minimize the Kullbak-Leibler Divergence between them through the gradient descent algorithm, normally training a neural network. By reducing the differences between both probability distribution, a high dimension space is transformed to a low dimension space, preserving the relationships among data entries given the values of its features.

For calculating the t-SNE, given a dataset X with a length N, for each pair of data entries xi and xj, the probability distribution in first stage is calculated by

$$P(i,j)=\frac{P\left( i | j \right)+ P(j|i)}{2N},$$

where the conditional probability$P\left( i | j \right)$ is computed by using

$$P\left( i | j \right)=\left\{ \begin{aligned} \frac{\exp\left( -\frac{\left\| x_{i}-x_{j} \right\|^{2}}{2\sigma_{i}^{2}} \right)}{\sum_{k\neq i} \exp\left( -\frac{\left\| x_{i}-x_{k} \right\|^{2}}{2\sigma_{i}^{2}} \right)}, &if i\neq j, \\ 0, &if i=j. \end{aligned} \right..$$

In last expression, $\left\| x_{i}-x_{j} \right\|$ denotes the Euclidian distance between data entries but another distance metric could be used. For second phase, points Y = (y_0_, y_1_, y_2_, …, y_N_) are found in such a way that minimize the Kullback-Leibler Divergence between $P(i|j)$ and $Q(i|j)$, where if $i\neq j$ then

$$Q(i|j)=\frac{\frac{1}{1+\left\| y_{i}-y_{j} \right\|^{2}}}{\sum_{k} \sum_{k\neq l} exp(\frac{1}{1+\left\| y_{k}-y_{l} \right\|^{2}})}$$

and it is equal to 0 in another case.

**Linear Projection for Visualization of multidimensional datasets**

Data Visualization is a powerful tool for data analysis in several stages of data processing. It plays a very important role in exploratory data analysis and data mining tasks. However, the visualization of datasets with more than two or three dimensions (variables) represents a challenge using conventional plotting 2D or 3D.

In order to provide a solution to this problem, a series of methods have been developed in which linear projections are made in order to reduce multidimensional data that can be presented in two or three dimensions. Several methods exist for performing that linear projections, examples are: Lineal Discriminant Analysis, Principal Components Analysis, Orthogonal Projection, Linear Regression, among others.

In general, a linear projection could be described as a linear transformation P from a vector space to the same vector space that satisfies P2 = P. For a given dataset several projections could be performed resulting in a variety of graphs expressing different facets of the dataset. When a class separation is the objective for the projection, choosing the appropriate parameters and method is crucial like could be appreciate in the image below.

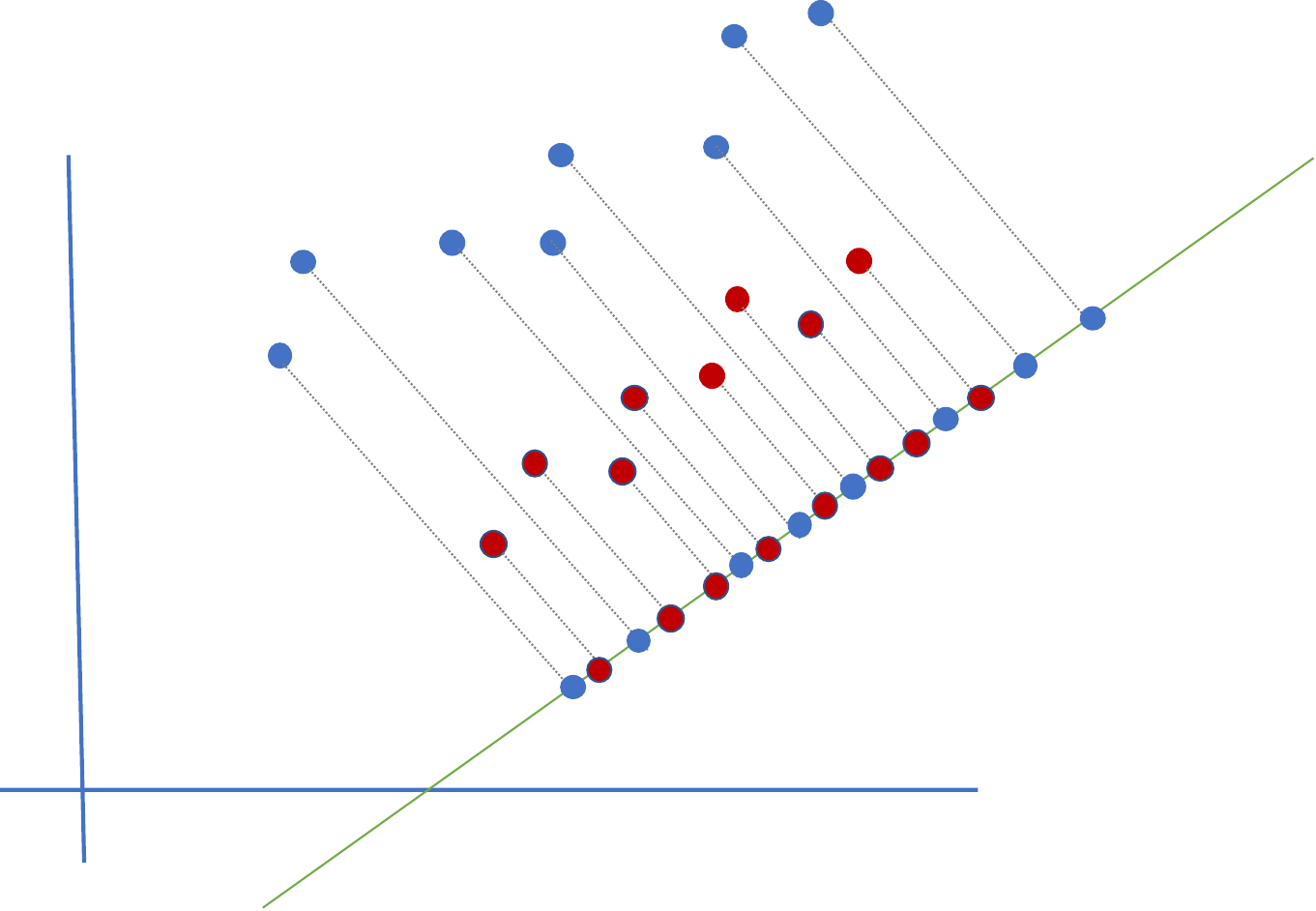

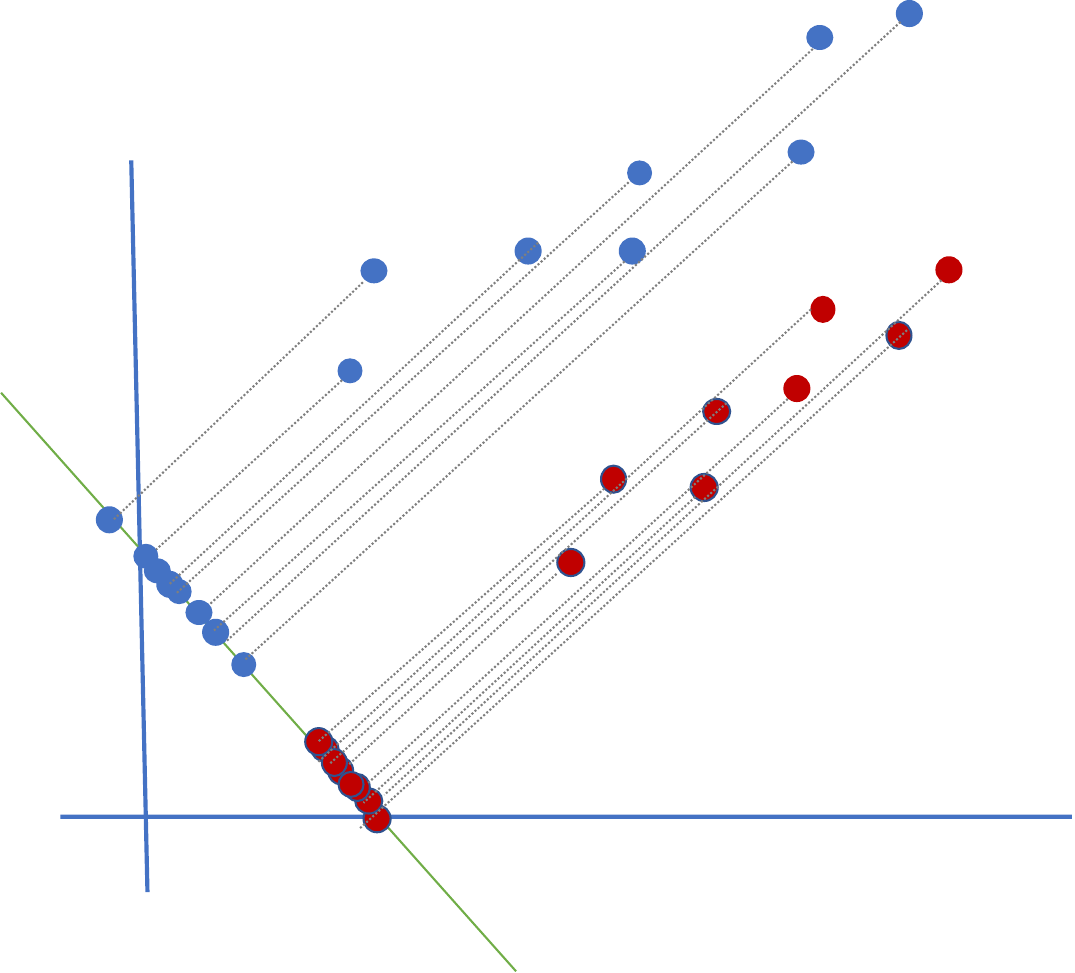

Fig. a	Fig. b

In the figure, we observe the same class-labeled multidimensional dataset being projected over two different linear models. In a) the resulting projection combines the entries from the two classes and does not help to separate them, while in b) the same dataset is correctly separated by classes, in both a linear model has been created for the projection, being different in the parameters used to construct them, showing the benefits obtained from an adequate linear projection for analyzing multidimensional datasets. In this work the Linear Projection was conducted using Orange© 3.26

**K-means**

K-means is an algorithm that aims to generate clusters of an observed dataset $\{x_{i}\}$ of n observations into k groups. Each group is represented by the average of the points that compose it. The centroid is the representative of each group. The number of groups to discover, *k*, is a parameter that must be configured a priori. The clustering method starts with *k* randomly located centroids and assigns each observation to the closest centroid. After being assigned, the centroids are moved to the average location of all assigned data and the points are assigned again according to the new positions of the centroids.

The objective of K-means is to group the observations in such a way that all those that are in the same group are the most similar to each other and that those that belong to different groups are the most dissimilar to each other. Distance measures, such as Euclidean, are used to measure similarities and differences. A measure of how well the centroids $M^{t}=\left( m_{1}^{t},m_{2}^{t},\ldots,m_{k}^{t} \right)$, at iteration $t$, represent the members of your group is the sum of the squared errors. K-means, in each iteration *t*, tries to reduce the value of the sum of the errors squared. The measure consists of the sum of the squared distances of each observation $x_{i}$from the centroid of its group

$$M^{t}=argmin\sum_{j=1}^{k} \sum_{i=1}^{n} ||x_{i}-m_{j}^{t-1}||.$$

The algorithm is iterative and always concludes, either by fixed number of iterations or when it converges to a solution (total error due to assignations remains the same than in the previous iteration). Since it does not necessarily find the most optimal configuration, the one that corresponds to the minimum of the objective function is shown. Finding a minimum of the function, even if it is not the absolute minimum, guarantees a grouping in which the groups are sparsely dispersed and separated from each other. The algorithm is significantly sensitive to centroids that are initially randomly selected. This effect can be reduced by performing multiple runs of the method. The clustering by K-means was conducted using Orange© 3.26
